# From Molecular Insight to Mesoscale Membrane Remodeling: Curvature Generation by Arginine-Rich Cell-Penetrating Peptides

**DOI:** 10.1101/2025.04.14.648709

**Authors:** Katarína L. Baxová, Jovi Koikkara, Christoph Allolio

**Affiliations:** Institute of Organic Chemistry and Biochemistry, Czech Academy of Sciences, Flemingovo nám. 542/2, 160 00 Prague, Czech Republic; Charles University, Faculty of Mathematics and Physics, Mathematical Institute, Sokolovská 83, 186 75 Prague 8, Czech Republic

**Keywords:** Multiscale modelling, Arginine Magic, Chemical specificity, Cell-penetrating peptides, Molecular dynamics simulations, Monte Carlo simulations

## Abstract

The enhanced cell penetration ability of arginine-rich peptides, such as nonaarginine (R_9_), compared to their lysine-rich counterparts remains incompletely understood. Atomistic simulations reveal that R_9_ binds significantly more strongly (≈ 20 kJ*/*mol) and penetrates deeper into anionic lipid headgroup region than its lysine equivalent. This enhanced interaction translates into a stronger induction of negative membrane curvature by R_9_. We combine these data to construct a model of peptide binding and curvature induction in fusogenic lipid mixtures and employ a multiscale simulation approach, combining atomistic molecular dynamics (MD) with mesoscopic Monte Carlo (MC) simulations, to dissect the molecular basis and morphological consequences of arginine specificity. The results show that stable membrane invaginations, as observed in studies of cell penetration, require excess membrane and are stable only for R_9_ (outside of the domain of stability of the Helfrich stomatocyte). By analyzing lipid and protein sorting coupled to the membrane structure, we explain the interplay of Gaussian and mean curvature in providing a mechanistic basis for the initial membrane deformation events potentially involved in ‘Arginine Magic’ cell entry pathways.

## 1 Introduction

Cell-penetrating peptides (CPPs) are an important means of drug delivery: They are able to transport large, polar cargo across the plasma membrane.^1^ *Arginine Magic* denotes the ability of arginine-rich CPPs to enter cells.^2^ The positively-charged peptide nonaarginine (R_9_) is an efficient CPP, in contrast to nonalysine (K_9_), which has the same charge. This (molecular) ion-specific effect has been studied on artificial phospholipid mixtures rich in phosphatidylethanolamine (PE) and phosphatidylserine (PS) as a model systems for cell penetration. ^3–5^ It was found that these model membrane systems selectively undergo fusion and form multilamellar structures^3^ as well as cubic phases upon R_9_ addition. ^4,5^ Analogous multilamellar structures have also been documented inside cells on multiple occasions.^3,6,7^

On this occasion, it should be pointed out, that the actual cellular entry mechanism was reported to be dependent on heparin binding receptors,^8^ that membrane proteins are necessary for passive entry even into giant plasma membrane vesicles (GPMVs)^9^ and that the artificial lipid compositions susceptible to leakage do not correspond to the composition of the plasma membrane.^3^ Nevertheless, the structural analogies of the PE and PS model system have held up well - even the experimentally known selectivity for arginine over lysine transfers to this model. Hence, model lipid bilayers have been extensively studied via molecular dynamics simulations.^10–13^

R_9_ has often been reported to induce membrane curvature,^3,4,14–16^ either positive, negative or negative Gaussian in nature. In recent years, there has been a forming consensus that membrane curvature induction is a key component of the cellular entry mechanism.^16^

Despite the recognized importance of curvature, the precise mechanism by which these peptides induce membrane curvature remains poorly understood, making further investigation crucial. Curvature generation by proteins has been a subject of intensive research already for several decades.^17–24^ Some of our work focused on quantitatively extracting membrane properties in the presence of proteins and adsorbates. ^25–27^ In particular, we made predictions based on examining negative curvature generation by cationic adsorbates on negatively charged lipids.^25,27^ These predictions also extend to pore formation. ^28^ In contrast to *in-vivo* processes, such as bacterial division,^29^ the relative simplicity of the model membrane system allows for the possibility of a thoroughly, albeit sparsely, parametrized multiscale model. Most recently, Pei and coworkers have postulated a revised mechanism of entry for cell-penetrating peptides, which departs from inward budding, presumably stabilized by *negative* membrane curvature.^7^

While previous studies have qualitatively observed R_9_-induced curvature or modeled peptide-lipid interactions, ^30^ a quantitative, multiscale understanding linking specific molecular binding events (R_9_ vs K_9_) to the resulting changes in membrane material properties (curvature, bending rigidity) and subsequent large-scale morphological transitions (such as budding or invagination) is lacking. Here, we revisit the molecular basis for *Argine Magic* in the domain of lipid binding, we quantify and explain the induction of curvature on pure phospholipid bilayers and incorporate the resulting data into a simple model for arginine binding on mixed model bilayers using large-scale molecular dynamics (MD) simulations and then incorporate it into our recently developed general Monte Carlo (MC) toolkit for membrane deformations, which couples it to the mesoscopic deformations.^31^ This enables us to investigate the conditions and physical determinants of inward budding as a proposed initial step of cell penetration.

## 2 Methods

### 2.1 Molecular Dynamics Simulations

A membrane bilayer containing 1024 lipids per leaflet was built by CHARMM-GUI^32^ containing DOPE:DOPS:DOPC lipids (60:20:20). 64 molecules of nonaarginine or nonalysine were used along with 50 TIP3P water molecules per lipid and 150mM KCl plus additional ions to counteract peptide charges. Both the traditional CHARMM36/LJ-PME and charge-scaled prosECCo75^33^ force field systems were set to run in Gromacs^34^ for 1 *µ*s while the last 200 ns were used for analysis.

Nosé-Hoover^35,36^ temperature coupling with a 1 ps time constant for the groups of protein, membrane and water with ions alongside with semi-isotropic Parrinello-Rahman^37^ pressure coupling (1 bar) ensured the isothermal-isobaric ensemble. The long-range interactions were treated by Particle-Mesh Ewald electrostatics^38^ with a cutoff of 1.2 nm. Hydrogen atom bonds were constrained by LINCS. ^39^ The timestep for the leap-frog integration algorithm was set to 2 fs.

#### 2.1.1 Material Properties

For the computation of elastic properties, simulations of pre-equilibrated bilayers were continued for 200 ns using Gromacs 2020.3.^34^ We simulated at 300 K temperature, using the same ensemble, algorithms and cutoffs as in the previous section. Long-range electrostatics were treated using the particle-mesh Ewald method.^38^ Since the pressure decomposition does not support the SETTLE algorithm,^40^ the triangular geometry of water was constrained using LINCS with order five.^39^ No dispersion corrections or potential switching were applied. In addition, we removed the CMAP terms, as we did not yet implement a force decomposition for these, and R_9_ and K_9_ are unstructured. The timestep was 2 fs. 20.000 MD snapshots with velocities were taken from each simulation.

The simulated systems were pure membranes, containing 14000 TIP3P water molecules and 128 lipids, in addition to 150 mM ions (and neutralizing counterions in the case of DOPS). When adding peptides, we added 14 R_9_ or K_9_ to DOPS and 6 R_9_ or K_9_ to DOPC and DOPE, respectively, for the simulation of a fully covered membrane. The simulations were run in the presence of 150 mM ions and counterions. For the stress tensor^27,41^ and ReSiS^42^ extraction simulations, we used the unscaled CHARMM36m. Further simulation details for stress-tensor computations and details for the free energy profiles are given in the Supporting Information.

### 2.2 Mesoscopic Monte Carlo Simulations

Monte Carlo (MC) simulations to sample the equilibrium shapes of vesicles (including thermal fluctuations using the Metropolis algorithm to achieve a canonical distribution) were performed using OrganL, ^31^ our Dynamically Triangulated Surface (DTS) code. The model implementation for this paper will be uploaded here.

The osmolyte concentration *c*_0_ was set to 300 mM (300 mOsm) in agreement with standard PBS buffers, and the area compressibility modulus *K*_*A*_ was set to 200 pN*/*nm.^43,44^ We initialized the system with a uniform lipid composition of DOPE:DOPC:DOPS 60:20:20 and a protein coverage of *ϕ*_*p*_ = 0.5, with lipid and protein binding parameters detailed in Table S1 of the Supporting Information (SI). At this peptide coverage (50%), all PS lipids are neutralized, along with a fraction of PE lipids, a loading chosen to align with molecular dynamics (MD) potential of mean force (PMF) calculations. The bound peptide-to-lipid (P/L) ratio is 1 : 46. While the total protein load per vesicle was not directly measured, we estimate it by combining the rapid saturation of R_9_ binding on supported bilayers, ^11^ our computed high adsorption energies, and the empirically low R_9_-R_9_ repulsion.

This simulation setup was designed to directly compare the stability of competing, potentially metastable morphologies, which we categorize broadly as “spheroid” and “stomatocyte”. Given that Helfrich energy minima are often metastable, we ensure a robust identification of the global minimum by independently evolving both stomatocyte-like and prolate spheroid-like geometries and systematically comparing their energies. See section 2.5 of SI for more details.

#### 2.2.1 Analysis

To quantify global parameters in MC simulations, a *thermal averaging approach* was used. Specifically, the final 500 snapshots of simulations, spanning the last 500k steps, were selected for analysis. To account for the mesh’s translational and rotational degrees of freedom, individual meshes were aligned using the Iterative Closest Point^45^ (ICP) point cloud registration algorithm. Local properties were computed as face-wise averages and mapped onto the corresponding faces of the ICP-generated mesh. Additionally, a *bin averaging* procedure was employed to analyse local parameters to preserve spatial correlations for curvature plots. See section 2.6 of the SI for more details.

## 3 Results and Discussion

### 3.1 Peptide Binding to Pure and Mixed Lipid Bilayers

#### 3.1.1 Pure Membranes

We computed potentials of mean force (PMF) of peptide binding to single-component lipid bilayers. The PMFs (Fig. 1) consistently show stronger binding of R_9_ over K_9_ to lipid membranes, in particular to the pure DOPS bilayer, where the difference is *>* 20 kJ*/*mol (see Table 1 for the binding energy values and errors for R_9_, while the binding energy of K_9_ on DOPS bilayer is −51.4 ± 2.3 kJ*/*mol). In particular, there is no significant binding of K_9_ to the neutral PC and PE bilayers, however, the repulsion is lower for DOPE. There also appears to be a slightly higher binding affinity to DOPE for R_9_. The difference between the charged DOPS and the other lipids is mainly due to the electrostatics of binding. We used the scaled-charge Prosecco forcefield for the computation of binding free energies. ^33^ Charge scaling reflects the implicit electronic contribution to the susceptibility of the medium. When comparing with similar previously published work,^30^ we estimate that our binding free energies are slightly lower than those expected when using an unscaled force field. Interestingly, the binding energy difference between K_9_ and R_9_ on PS is larger than for the other bilayers, hinting at some specific preferential interaction of R_9_ with the PS headgroup.

**Table 1:** MD data for protein-lipid bilayer interactions. Γ and *g*_ext_ are the excess number of lipids per peptide and the extremum of the first RDF peak in Fig 4, respectively. Errors inΓ and *g*_ext_ are the absolute differences between the mean (averaged over “Prosecco” and “Charmm”) and individual model values.

| $R_9$ | $\Gamma$ | $g_{\text{ext}}(r)$ | $\Delta G_{\text{bind}}$ [kJ/mol] |
| --- | --- | --- | --- |
| DOPE | $0.3 \pm 0.6$ | $0.8 \pm 0.02$ | $-23.2 \pm 2.6$ |
| DOPS | $1.3 \pm 0.7$ | $1.48 \pm 0.10$ | $-86.0 \pm 4.3$ |
| DOPC | $-1.7 \pm 1.4$ | $0.54 \pm 0.13$ | $-19.6 \pm 3.0$ |
| Mixture (PE:PC:PS 0.6:0.2:0.2) | | | $\Delta G_{\text{bind}}$ [kJ/mol] |
| $R_9$ | | | $-50.0 \pm 1.4$ |
| $K_9$ | | | $-23.3 \pm 0.8$ |

**Figure 1:**
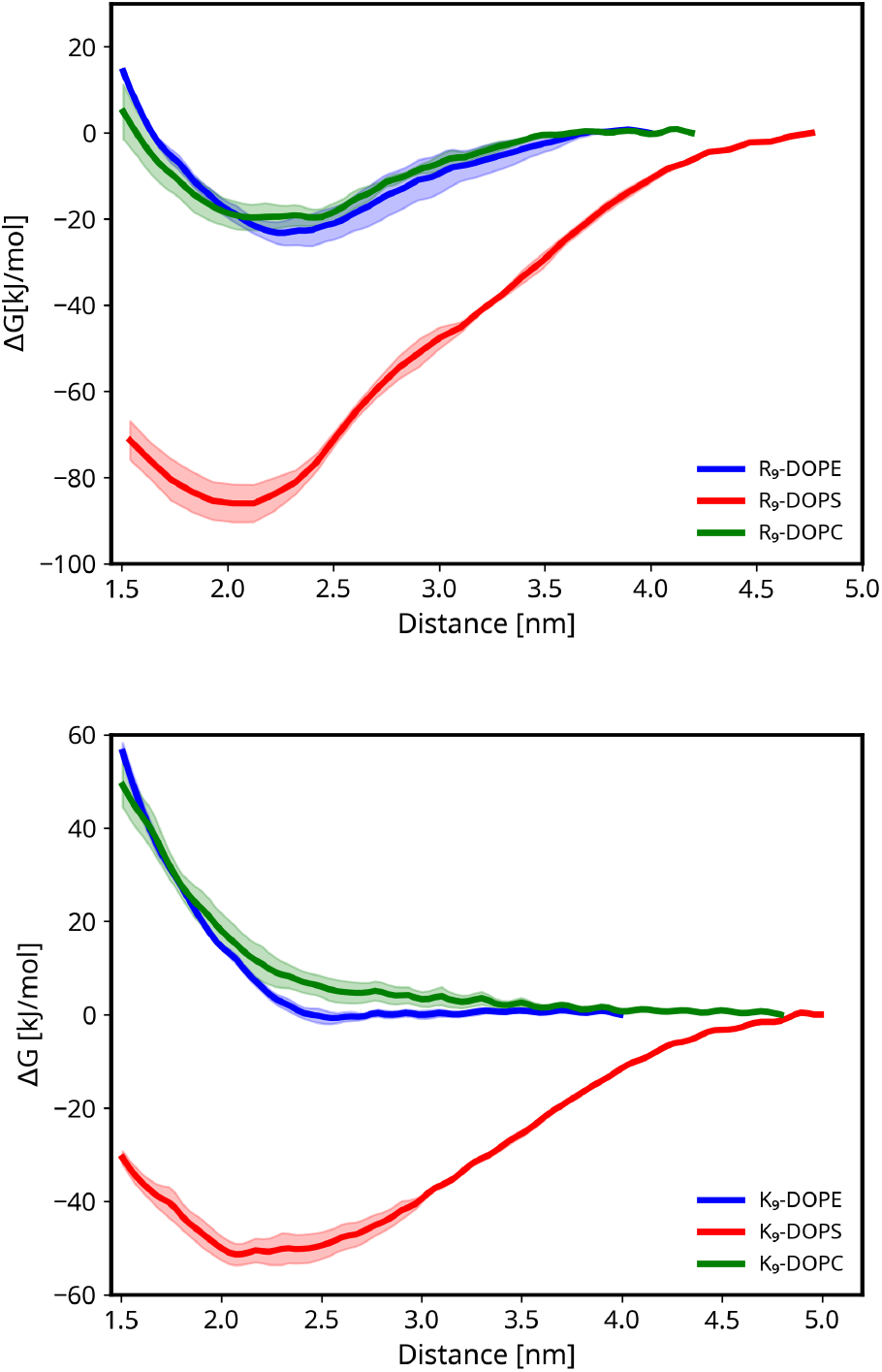
Top panel: Free energy profile of a single R_9_ binding to a lipid patch composed of DOPE (blue), DOPS (red), DOPC (green). Bottom panel: equivalent energy profiles for membranes interacting with K_9_.

These binding free energies show enhanced binding of R_9_ over K_9_ to all simulated lipid membranes. The long-range electrostatics do not differ (due to the identical sidechain charge). This implies that if the binding energy is part of the “magic”, it must lie in the stronger binding of arginine sidechains to membranes.

#### 3.1.2 Mixed Membranes

We also report binding energies of the peptides to mixed membranes in Table 1. As expected, binding to the mixtures is stronger than to pure neutral lipids but weaker than to the charged pure DOPS. To examine the structural factors of this binding in a general setting, we use results from our large-scale mixed-membrane simulation. The membrane contains all examined lipids at the ratios DOPE/DOPS/DOPC 0.6:0.2:0.2. Analyzing this simulation enables us to disentangle the molecular picture at the interface. In Fig. 2, we compare the density profiles of different species at the membrane interface in the presence of the peptides. The figure also compares scaled Prosecco and unscaled CHARMM36 forcefields.

**Figure 2:**
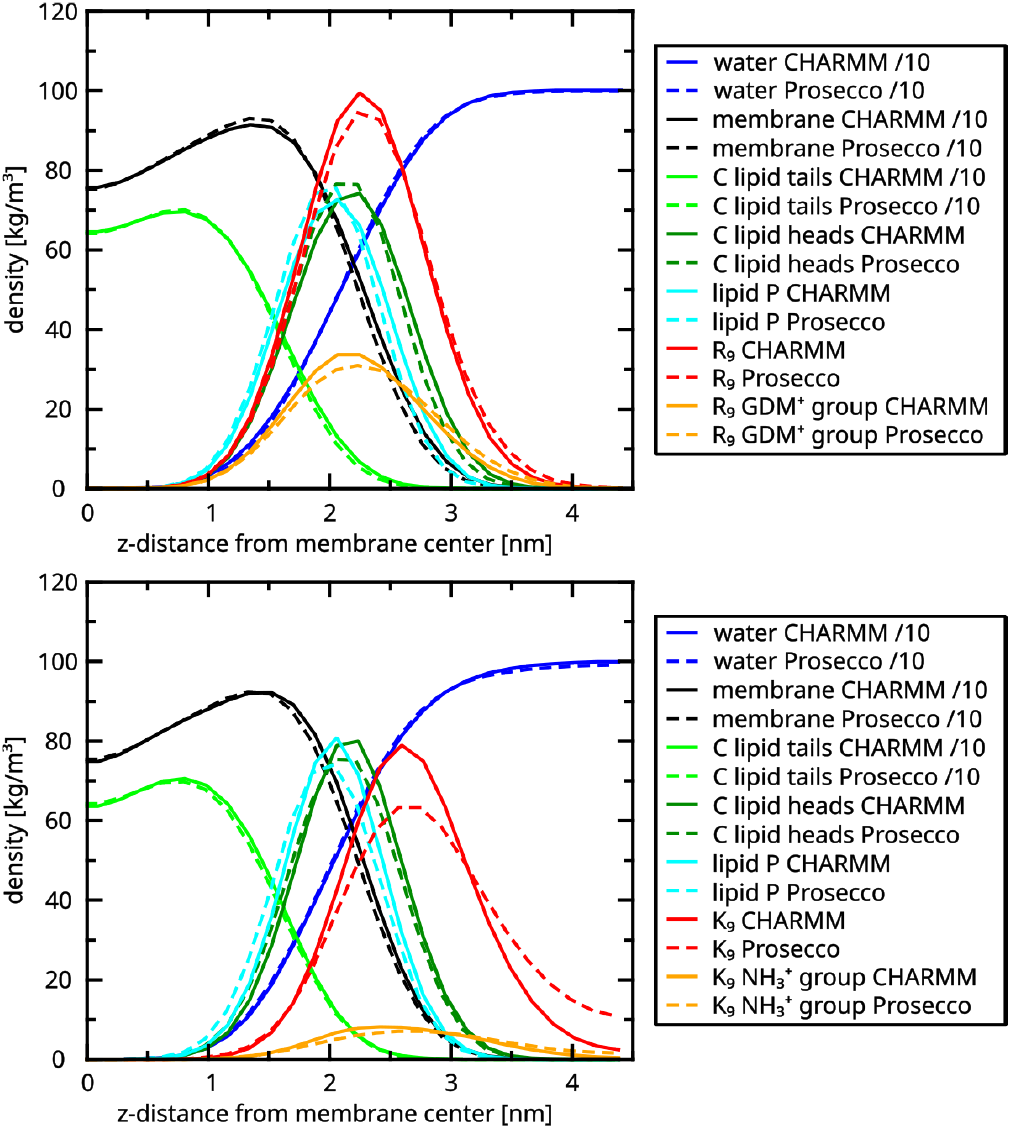
Density profiles from the large-scale simulations of R_9_ (top) and K_9_ (bottom) centered on the membrane midplane, showing both CHARMM (full lines) and Prosecco (dashed lines) results. Density profiles of all membrane atoms (black), water (dark blue), lipid tail C-atoms (green), phosphate P atoms (light blue), proteins (red) and headgroups (orange). Density profiles of water and lipids are scaled by 0.1 for clarity.

The striking difference between R_9_ and K_9_ lies in the far deeper binding of the arginine sidechain inside the membrane (yellow, upper part) compared to the lysine sidechains (yellow, lower part). The arginine sidechain density is collocated with the headgroup/phosphate position, whereas lysine is mainly distributed above the headgroup area. The deeper penetration of the arginine sidechain brings it closer to the locus of negative charge. This enhances the electrostatic interaction energy as well as the mutual screening of the charges. The picture is similar to the hydrophobic insertion mechanism by Kozlov et al., ^22^ except that in this case, the interaction is ionic and not hydrophobic and the steric component is minor. Yet, it is widely known that arginine is more hydrophobic than lysine, and we believe this is part of the explanation for why arginine is able to enter membranes more deeply.

#### 3.1.3 Arginine-Arginine Interaction

A crucial question regarding R_9_ behavior is whether R_9_ peptides interact strongly with each other on the membrane surface. Past studies showed that guanidinium moieties of the arginine sidechain do not repel in aqueous solution and even interact weakly. ^10^ In addition, aggregation of R_9_ on membranes has been observed in simulation^12^ and recently also in experiment.^46^ In contrast to this, the binding of R_9_ to membranes charged at 20% was previously found not to be cooperative. ^11^

To test this again, in the context of our lipid mixture, we computed two-dimensional radial distribution functions (RDF)s of a large number of R_9_ molecules on a flat lipid patch (see Methods). The results are shown in Fig. 3. These data are not consistent with a strong binding between poly-R_9_ molecules, as the RDF does not exhibit major peaks but fluctuates around unity away from a depletion/exclusion zone. It seems reasonable to approximate this behavior by a simple excluded volume of each R_9_, in agreement with the data by Cremer.^11^ In contrast to R_9_, K_9_ molecules were found by Cremer to repel on the membrane significantly. Recent experiments, ^46^ suggesting aggregation by R_9_ on membranes, do not furnish evidence of more than a stochastic collocation. Therefore, we suggest treating R_9_ chains as noninteracting on (negatively charged) lipid membranes.

**Figure 3:**
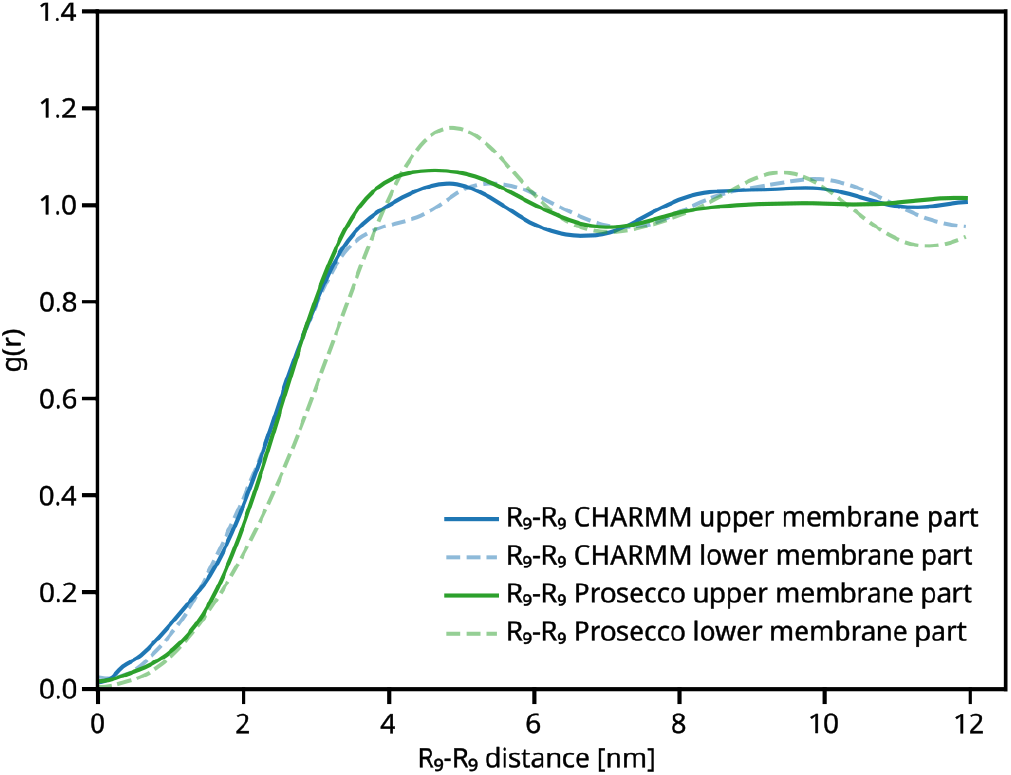
Arginine aggregation: Two-dimensional RDF of R_9_-R_9_ in the upper (solid lines) and lower (dashed lines) leaflet of the large membrane simulation.

#### 3.1.4 Arginine-Lipid Demixing

Having established the weak mutual-interaction of R_9_ peptides, we now turn to its interaction with lipids and the crucial phenomenon of lipid demixing. We already found that R_9_ strongly interacts with lipids, depending on the lipid headgroup type, in particular its charge. The long-term large-scale MD simulations of the membrane with peptides allow us to compute a realistic two-dimensional RDFs of the lipid molecular centers of mass vs R_9_ at the membrane surface in Fig. 4. The strong binding of R_9_ to negatively charged PS lipids is enough to generate a significant amount of demixing This is evidenced by a high peak of R_9_ under the protein accompanied with depletion of the other lipids. This depletion is larger for DOPC than for DOPE, as observed in both charge-scaled and unscaled simulations. It suggests a strong R_9_ preference towards PS at the expense of PC local lipid density, while PE remains almost unaffected. The demixing can be quantified both in coordination numbers and in excess quantities. We report estimates on an excess number of lipids per peptide (Kirkwood-Buff integrals) and the peak of RDF, together with the binding energies in Table 1.

**Figure 4:**
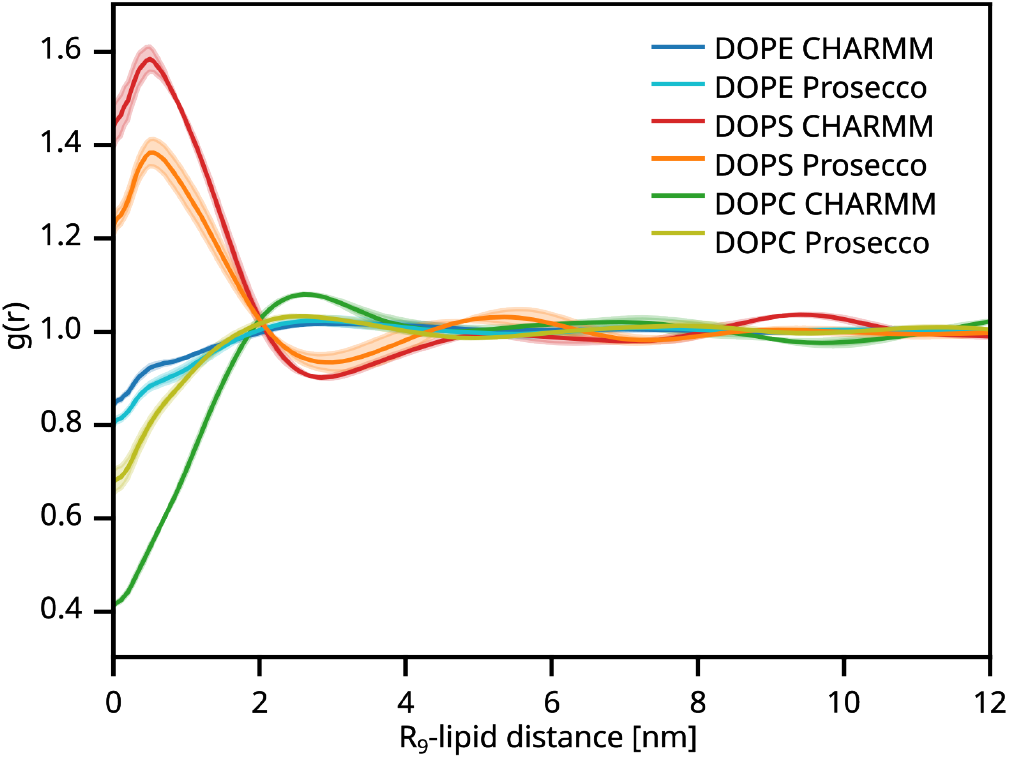
2D-RDFs of lipids around R_9_: R_9_-DOPE RDF in blue and cyan, from CHARMM36 force field and (scaled) Prosecco simulations, respectively; R_9_-DOPS RDF in red and orange and R9-DOPC RDF in blue and cyan.

The high peptide concentration in these big leaflet simulations was chosen to improve sampling, as lipids will not have to diffuse far to bind to a peptide. It, however, also means that of the 32 lipids for each protein, a significant amount will be in direct contact. This naturally limits the amount of lipid demixing, so the amount of lipid selectivity is probably not as high as in lower concentrations.

Our explicit atomistic simulation results can be compared with an earlier study by Harries et al. using a Poisson-Boltzmann/ideal mixing-based model.^47^ They found only weak demixing of PS in proximity of a single adsorbed K_13_ molecule. The highest increase in PS was a factor of 1.5, which is not that different from our observed RDF peaks. Note that the PB model ignores the interfacial “burying” of R_9_ to the membrane headgroups.

### 3.2 Peptide Binding Effect on Material Properties

We simulated a representative set of bilayers using the unscaled CHARMM model and computed the bending moduli *κ* and a bilayer tilt moduli 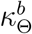 with the ReSIS method, which is based on local tilt fluctuations.^42,48^ We previously computed these values for DOPE, DOPC and DOPS lipid membranes, but recomputed the value for DOPS to show reproducibility of previously published simulations.^27^ To examine the influence on membrane bilayer properties, we put the membrane in contact with high (charge neutralizing) loads of R_9_, which we estimate to be close to saturation.

#### 3.2.1 Material Properties

To understand how R_9_ and K_9_ modify the material properties of lipid bilayers, we performed simulations on a representation set of bilayers. We used 6 R_9_ for uncharged lipids and 14 peptides for charged membranes to achieve approximate charge neutrality. The results, including equilibrium areas per lipid *A*_0_, are in Table 2. Since K_9_ does not bind to the uncharged lipids, as per our PMFs, we did not simulate the corresponding membranes. The influence of the peptides on elastic properties is not large for either peptide. However, it appears that K_9_ stiffens DOPS membranes, while R_9_ might slightly soften them. This effect is also visible in the area per lipid and might be attributed to a surface tension contribution of K_9_, which is offset by increased chain-packing. In contrast, the entry of R_9_ into the headgroup region might counteract some of the compressive effects of charge screening.

**Table 2:** Simulation results for bilayer mechanical properties. Bending moduli *κ* and spontaneous curvatures *J*_*s*_ are monolayer quantities. Areas per lipid *A*_0_ and tilt moduli 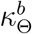 are bilayer properties. Superscript ^*a*^ from^25^ unscaled simulations at 303.15K. Errors are standard errors.

| System | $\kappa$ [kT] | $J_s$ [nm<sup>-1</sup>] | $A_0$ [Å<sup>2</sup>] | $\kappa_\theta^b$ [kT/nm<sup>2</sup>] |
| --- | --- | --- | --- | --- |
| DOPE<sup>a</sup> | 15.83 | -0.24 | 61.64 ± 0.16 | 14.76 |
| DOPC<sup>a</sup> | 11.56 | 0.0 | 68.02 ± 0.08 | 22.68 |
| DOPS | 14.27 ± 0.84 | -0.06 ± 0.01 | 63.38 ± 0.23 | 19.76 ± 1.23 |
| DOPE + 6 R<sub>9</sub> | 16.20 ± 0.80 | -0.30 ± 0.03 | 61.09 ± 0.14 | 23.15 ± 0.94 |
| DOPC + 6 R<sub>9</sub> | 10.88 ± 0.49 | -0.03 ± 0.02 | 68.60 ± 0.17 | 14.31 ± 0.56 |
| DOPS + 14 R<sub>9</sub> | 13.16 ± 0.60 | -0.26 ± 0.08 | 63.23 ± 0.14 | 19.48 ± 0.96 |
| DOPE + 6 K<sub>9</sub> | not binding acc. to PMF |  |  |  |
| DOPC + 6 K<sub>9</sub> | not binding acc. to PMF |  |  |  |
| DOPS + 14 K<sub>9</sub> | 16.26 ± 0.52 | -0.17 ± 0.02 | 61.49 ± 0.40 | 23.71 ± 0.84 |

#### 3.2.2 Stress Profiles

To gain deeper mechanistic insight into the curvature-inducing effects of R_9_ and K_9_, we computed local stress profiles for single-component lipid membranes. The largest impact on curvature is found for the pure DOPS membrane. Figure 5 illustrates the different effects of R_9_ and K_9_ on charged lipids. We interpret the peak around 2 nm as associated with headgroup repulsion. It is lower for R_9_ than for K_9_. The whole profile is damped for R_9_, potentially also due to packing effects, but the decay of the positive electrostatic pressure in particular is faster for R_9_. The positive pressure is due to the electrostatic repulsion of the adsorbed proteins. We attribute the lower repulsion both to the electrostatic screening by headgroup binding and, to some extent, to the compactness of the peptides due to the attraction between sidechains.

**Figure 5:**
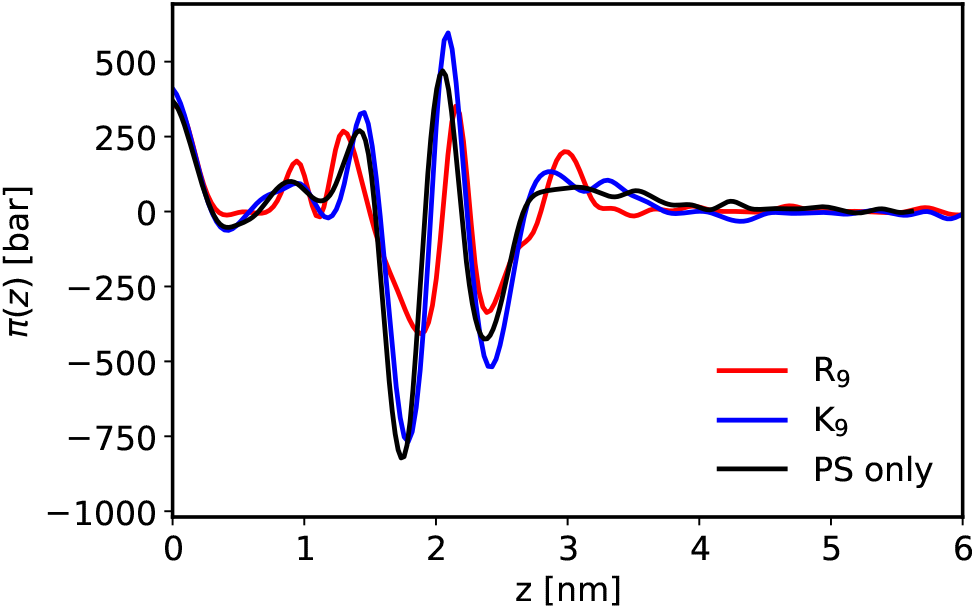
Comparison of the symmetrized lateral stress profiles of DOPS in the presence of R_9_ (red) and K_9_ (blue) and in the absence of peptides (black). On the *x* axis, *z* denotes the distance from the membrane center projected on the surface normal.

#### 3.2.3 Curvature Generation

By combining the computed stress profiles with the elastic properties, we can now quantitatively assess the curvature-generating potential of R_9_ and K_9_ using the established relationship^49,50^ between the first bending moment and the product of spontaneous curvature, *J*_s_ and *κ*:

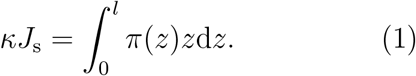

Here, integration is carried out along the membrane normal **ê**_*z*_.

A large negative curvature generating effect is observed for R_9_ on DOPS (see *J*_*s*_ in Table 2). This result is similar to what was observed in the case of Ca^2+^ ions^27^ on the same lipid. In our previous study, Ca^2+^ was shown to stabilize DOPS membrane fusion stalks as indicative of fusion. Note also that Ca^2+^ experimental behavior was similar to that of CPPs in this system. ^3^ The effect of R_9_ is significantly stronger than that at the same loading of K_9_. In particular, the increased *κ* on K_9_ reduces the effect on *J*_*s*_. It is remarkable that R_9_ can reduce the spontaneous curvature of uncharged DOPE. Our previous research found pure DOPE/DOPS to be very susceptible to fusion by R_9_ (over K_9_), and this effect was reduced by adding DOPC.^3^

The error bar values in the curvature calculations (Table 2) are standard errors, including error propagation (see Supporting Information). The J_*s*_ values given here were calculated automatically by integration over the whole box, which is sometimes conducive to errors as fluctuations in the pressure far from the membrane center can drastically influence the first part of the distribution.

### 3.3 Effective Mesoscopic Model

In order to understand the mesoscopic structural effects of the peptide and lipid specific interactions, we propose an effective continuum model. The basis of our model is the Helfrich-Canham-Evans theory. ^51–53^ This thin-shell theoretical model is a well-established way to describe membrane curvature elasticity and structure. Instead of constraints for area and volume, we introduce laterally compressible membranes and an osmotic pressure,^54^ as well as regular solution type mixing terms. The total energy of the system is then given by

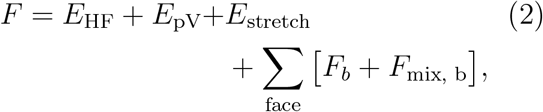

where

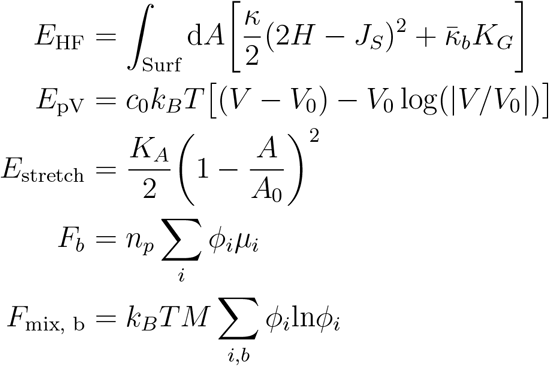

Here, the parameters for (Gaussian) bending rigidities 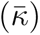 *κ* and spontaneous curvature *J*_*S*_ vary over the mesh but are constant on each face. Integration is performed on the faces. *H* denotes the mean curvature. The bilayer binding energy of the peptide *F*_*b*_ and the mixing energy *F*_mix, b_ are calculated per face and added. The volume work *E*_pV_ is calculated using the vesicle volume *V* from work against the osmotic pressure difference between the interior and exterior of the vesicle (assuming balanced osmolarity at V_0_) at a solution osmolarity *c*_0_. The area compression energy *E*_stretch_ is calculated from the total area *A* using the bulk modulus *K*_*A*_ and the stress-free reference area *A*_0_. *n*_*p*_, *ϕ*_*i*_ and *µ*_*i*_ are the local (mesh element) numbers of peptides, fractional peptide coverage and binding energy, respectively. See the Supporting Information for exact parameter values and mixing rules. We use a Metropolis Monte Carlo approach to sample the Boltzmann distribution of lipid film structures. For details on the numerical evaluation, lipid mixing dynamics and mesh evolution see. ^31^

#### 3.3.1 Model Assumptions

We assume the membrane to be a curvature elastic thin film, obeying the Helfrich theory; ideal lipid mixing and osmotic pressure expressions apply. The area per lipid is assumed to be unaffected by the lipid mixing. Proteins are shapeless and interact only via size exclusion and by covering lipids. Proteins modify properties only of those lipids that they “cover” (discussed below and in Section 3.1.3). Furthermore, long-range electrostatic as well as tilt deformation are neglected. We also neglect inertia so that all motion is due to thermal fluctuations. In our discretization, a protein will always cover exactly one triangle, there are no fractional occupations. More than one protein on each face is not allowed, as we do not know how to interpolate between populations. In contrast, in the case of lipid mixing, we use well-established mixing terms and fractional occupations. ^25,55^ These are described in detail in Section 1 of SI. These terms neglect potential differential stress and membrane asymmetry effects.^56,57^

#### 3.3.2 Model Parameters

Our MD simulations provide us with parameters for single-component bilayers, given in Table 2. Values for *A*_0_, *J*_*s*_ and *κ* for pure lipids and those in contact with the peptides from this table. We use an estimate from^27^ for the Gaussian bending modulus of the bilayer (see SI), as monolayer values are difficult to obtain.^58^

For the binding free energies *µ*_*i*_ of the lipids to the peptides, we utilize values within the error bars of our single-component PMF data. All model parameters are summarized in Table S1 of the SI. By using the extracted binding free energies and material properties from the MD simulations, we transfer the essential information into an effective model to examine the morphological consequences of the local changes in curvature-elastic parameters.

#### 3.3.3 Model Validation

Consider a flat membrane divided into two virtual membrane systems: one functionalized with a peptide and the other unmodified. The free energy concerning lipid and protein is,

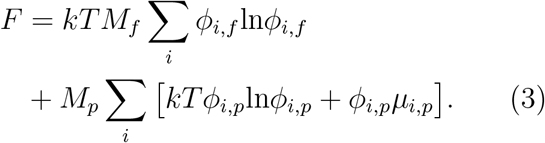

Since the membrane is flat, the bending terms will not affect the mixing. We also approximate ∀*i, j* : *A*_*i*_ = *A*_*j*_ ⇒ *ϕ*_*i*_ = *x*_*i*_, so that we can express the energy in mole fractions *x*_*i*_. We divide a lipid population into a free fraction *M*_*f*_ and a lipid population *M*_*p*_ bound to the protein with a binding energy *µ*_*p*,*i*_. The model Hamiltonian can, therefore, be expressed as:

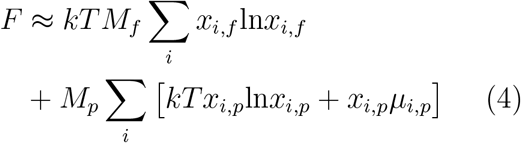

We also apply the following constraints:

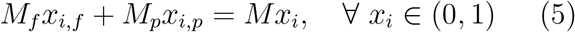

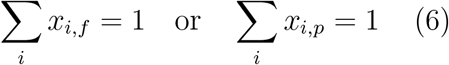

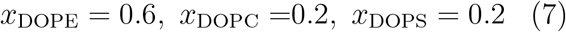

where *M* := *M*_*f*_ + *M*_*p*_ is the total number of lipids and *x*_*i*_ is the mole fraction of *i*th lipid, both of which are fixed. Applying either condition in the Eq. (6), in conjunction with Eq. (5), inherently satisfies the complementary constraint. The resulting energy is minimized against Eq. 4 for a given *M*_*p*_. Since the *M*_*p*_ value depends on the unknown peptide area, *M*_*p*_ is an adjustable parameter. We optimize it by minimizing a ℒ^2^ norm between the MD data (Table 1) and the estimated {*x*_*i*,*p*_} after converting the latter to excess quantities. The technical details are in SI.

For the large simulation with 1024 lipids per leaflet and 34 R_9_ bound to the leaflet, the optimal area of R_9_ is 14.786 nm^2^ (23.126 lipids per R_9_ and weighted average APL from Table 2 of SI) resulting in the excess number of lipids per peptide,Γ, as:

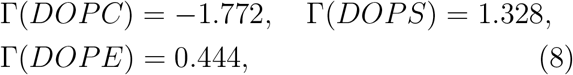

Here, the excess lipid number per peptide,Γ, is defined as the difference between the number of bound lipids after optimization and the initial number, which is uniformly distributed according to Eq. 7.

We find that Eq. 8 is in good agreement with Table 1. To give an idea of the molecular dimension, this area corresponds to a radius of ca. 2.2 nm, not too far from the RDF data.

Next, we consider a small system of 64 lipids and only one peptide. In this case, the coverage ratio is only 32.8%. For such a system, we predict

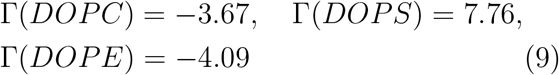

and a binding energy of 56.69 kJ*/*mol, in reasonable agreement with the MD data Table 1. Please note that in this system, demixing is predicted to be stronger than in the high peptide concentration large-scale simulation. This leads to strong binding. In this system, Monolayer curvature is predicted to be *J*_*S*_ = −0.25 (−0.17) nm^*−*1^ for the bound (unbound) zone, while monolayer *κ* = 14.16 (14.28) kT respectively. For K_9_, we use the respective *µ*_*i*_ from Table 1 and the identical area as for R_9_. Then the K_9_ binding energy prediction for the small patch is −27.7 kJ*/*mol, which is in good agreement with Table 1. Note that as K_9_ has a lower binding energy and has experimentally anti-cooperative behavior, ^11^ the model for K_9_ should be considered more of an upper bound of the effect, at least at identical peptide load. Our simplified model is able to capture key aspects of peptide-lipid interactions observed in atomistic MD simulations: It reproduces preferential binding, binding free energies on lipid mixtures, and the demixing under the peptides. We use the area prediction of R_9_ as input for our simulation. By its constant lipid coverage assumption, it does not describe well the fact that the protein coverage can be higher for strongly negatively charged membranes. This is to be expected for a model that does not explicitly treat electrostatics.

### 3.4 Mesoscopic Consequences of R_9_ Specificity

#### 3.4.1 Stability and Morphology

The equilibrium shapes of vesicles governed by the Helfrich energy with uniform *J*_*s*_ have been extensively studied. ^31,56,59^ Energetically stable shapes include stomatocytes, discocytes as well as prolate, and oblate spheroids. Helfrich minima are usually plotted as a function of the reduced volume:

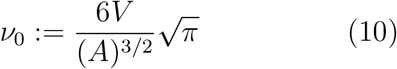

In the standard formulation, the reduced volume is computed from the total membrane area *A* and the enclosed volume *V* . It reflects the scale invariance of the Helfrich theory in the absence of spontaneous curvature. Thus, volume and area constraints are unified into one parameter. In our case, we do not have strict volume or area preservation - the values of *ν*_0_ correspond to the osmotic pressure difference minima of the free energy (Eq. 2), i.e. zero resultant osmotic pressure (*V*_0_) and the stress-free area (*A*_0_). As *ν*_0_ is linear in the volume, it can be interpreted as a degree of *filling* of the vesicle. It should be emphasized that the actual values of the instantaneous mesh are different due to, e.g. the mechanical forces working against volume preservation.

In the absence of spontaneous curvature, prolate configurations remain stable up to *ν*_0_ ≈0.65, while stomatocytes are global free energy minima for *ν*_0_ ∼0.55. In between these two values, the discocyte structure is stable. In our case, the system is not *a priori* scale-invariant as the individual lipids have defined intrinsic curvatures. Hence, the results are, in principle, valid only for our choice of vesicle size and osmolarity. Nevertheless, plotting as a function of *ν*_0_ illustrates the difference to the standard Helfrich results. Here, we investigate the stabilization of competing structures in the presence of lipids and peptides, with a particular focus on the “stomatocyte” shape. This geometry is of primary interest as it represents inward budding, a process hypothesized to be the initial step of CPP entry. ^7^

We compare its relative energy to that of prolate/oblate spheroid-type structures in Fig. 7. As we perform Monte Carlo simulations sampling the Boltzmann distribution, the “metastable”, i.e. higher energy state is inherently transitory: The underlying stochastic process will converge to the stationary distribution, which knows these “metastable” states only as fluctuations. Yet, these shapes persist for long enough to compute average energies, allowing for an estimate of the relative stability. To illustrate the transitory nature of unstable shapes, they are marked in grey.

**Figure 6:**
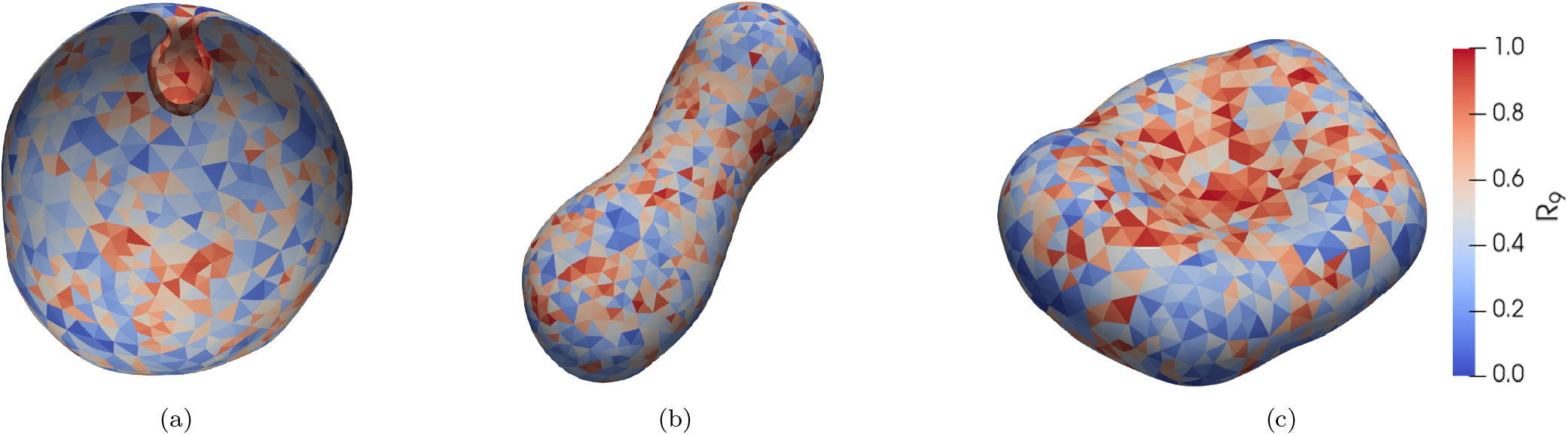
Representative structures with thermally averaged R_9_ occupancy. (a) Transverse cross section of stomatocyte of *ν*_0_ = 1.00, (b) prolate of *ν*_0_ = 0.75 and (c) discocyte of *ν*_0_ = 0.80.

**Figure 7:**
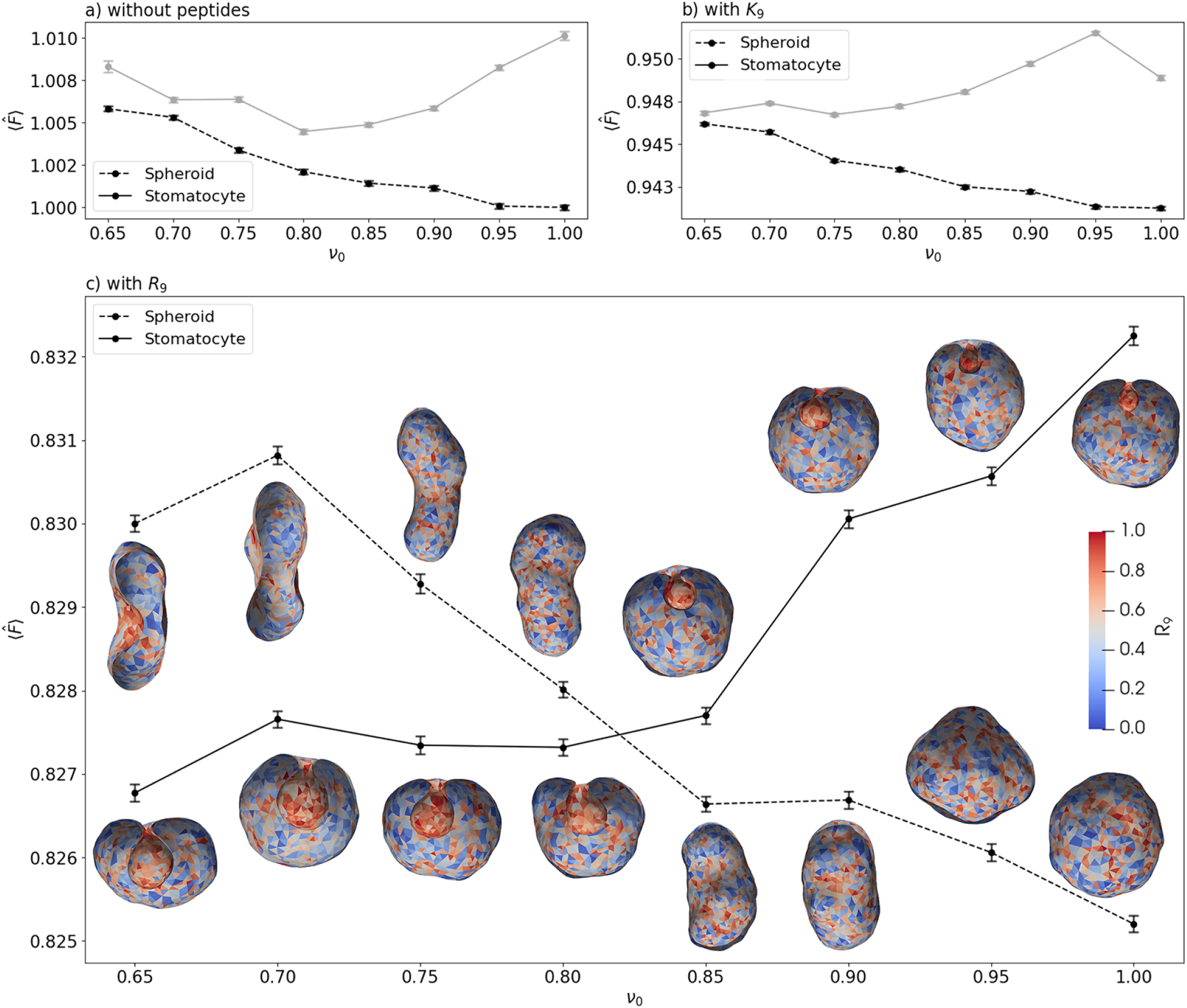
Model Free Energies. All energies are given relative to that of the spherical DOPE/DOPC/DOPS system at *ν*_0_ = 1.0 without peptides. Relative stability of spheroids vs stomatocytes are given: (a) without peptides, (b) with K_9_, and (c) with R_9_. Simulation snapshots depict the corresponding reduced volume, with color maps indicating average sorting occupancy. Grey plots denote metastable states. Error bars represent the SEM with 95% CI.

For example, in the control simulation without peptides (Fig. 7a), the stomatocyte shapes slowly transition towards prolate geometries. This behavior mirrors the predicted global minimum resulting from Helfrich energy minimization without lipids. In the absence of lipid demixing, there will be no spontaneous curvature on the mesh, as the intrinsic curvatures of monolayers of identical composition cancel. Hence, our result is in line with the established understanding within the community that ideal mixing entropy prevents lipid demixing to a large extent. Consequently, spatial curvature variations that stabilize curved membrane structures beyond the known Helfrich shapes require additional stabilizing mechanisms.^60,61^

The energy profiles of membranes containing K_9_ peptides (Fig. 7b) are similar to the control run in that *no stable* range for the stomatocyte structure is observed for *ν*_0_ *>* 0.65.

The presence of R_9_ peptides (Fig. 7c) significantly enhances the stability of invaginated structures compared to (prolate) spheroids. Energetically, invaginated and prolate shapes were comparable within a reduced volume (*ν*_0_) range of 0.80 − 0.85. Below *ν*_0_ ≈ 0.80, a clear divergence of energy profiles is observed, with stomatocyte geometries having considerably lower energy than spheroids, in contrast to what is observed in the control run or the simulation with K_9_. This is further evident in the Helfrich Energy (Eq. 2) contribution in Fig. 8, which shows a similar pattern suggesting the active role of spontaneous curvature generation. This plot also reveals that the invagination stability decreases with a reduction in excess membrane area, as osmotic pressure progressively destabilizes this geometry. A detailed energy decomposition plot is provided in Fig. S1 of the Supporting Information.

**Figure 8:**
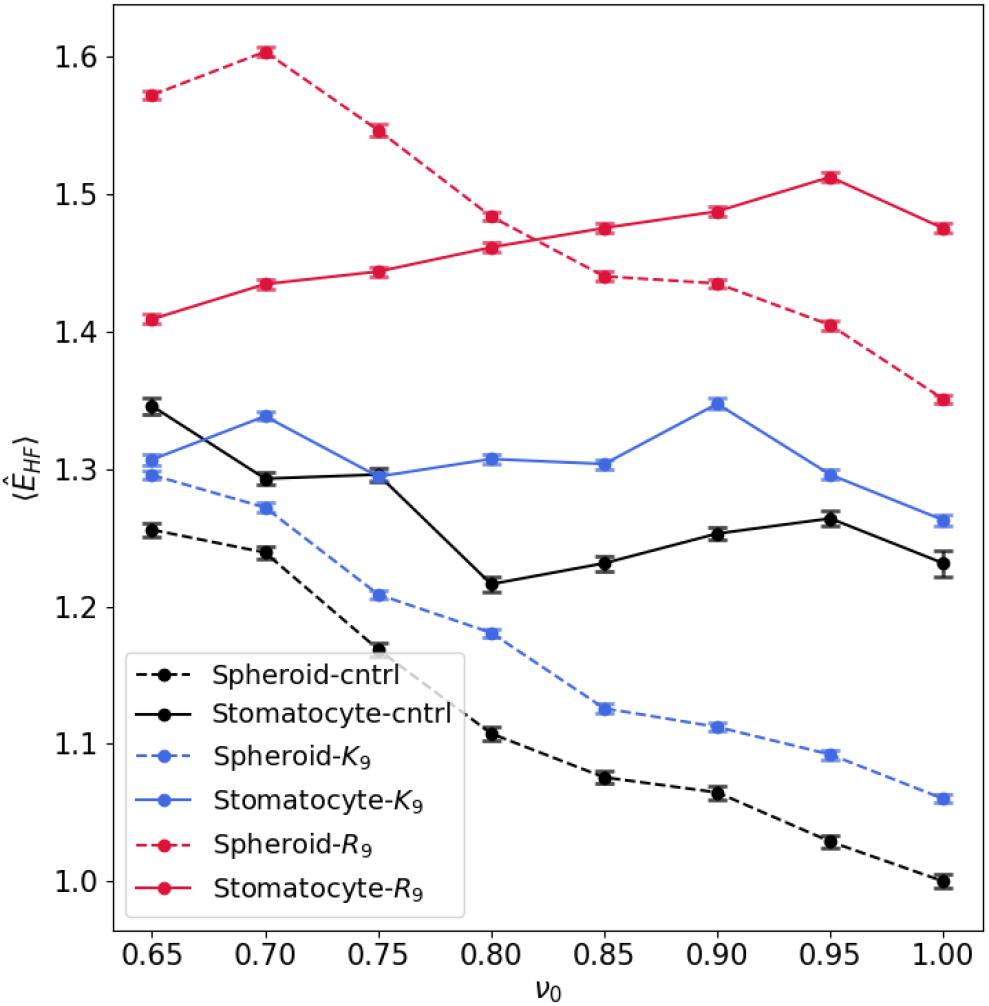
Contributions of curvature elastic term (Helfrich Energies). Profiles are shown under three conditions: control (black), with K_9_ (blue), and with R_9_ (red). Dashed lines represent spheroid morphologies, while solid lines indicate stomatocyte morphologies. Error bars denote the SEM with 95% CI. All energies are given relative to that of the spherical DOPE/DOPC/DOPS system at *ν*_0_ = 1.0 without peptides.

Collectively, these findings demonstrate that R_9_ peptides stabilize inward budding in DOPE/DOPC/DOPS vesicles, requiring a minimum excess area or hyperosmotic stress for stable invagination formation. We attribute this finding to the spontaneous curvature generation by R_9_, which must be concomitant with significant lipid demixing.

#### 3.4.2 Curvature Sorting

We investigate the underlying mechanisms by examining the spatial organization of R_9_ and lipids in representative geometries. In order to quantify curvature sorting, we define the *fractional excess coverage* as

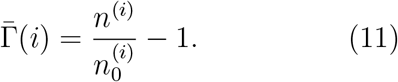

Here, *n*^(*i*)^ is the occupancy (number of lipids) on a face 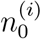 being the equivalent of the *i*^th^ compound in the initial uniform distribution. Figure 9 displays heatmaps of the fractional excess R_9_ coverage on a fixed mesh representing a typical “stomatocyte” or inward-budded vesicle with *ν*_0_ = 0.80, calculated by first binning the instantaneous mesh data to preserve spatial correlations and averaging over the last 200 such snapshots. The fixed mesh was employed to enhance the convergence and isolate the sorting effect. However, the findings remain consistent, albeit with reduced prominence, when using an unfrozen mesh and are included in the SI Fig. 2 for completeness.

**Figure 9:**
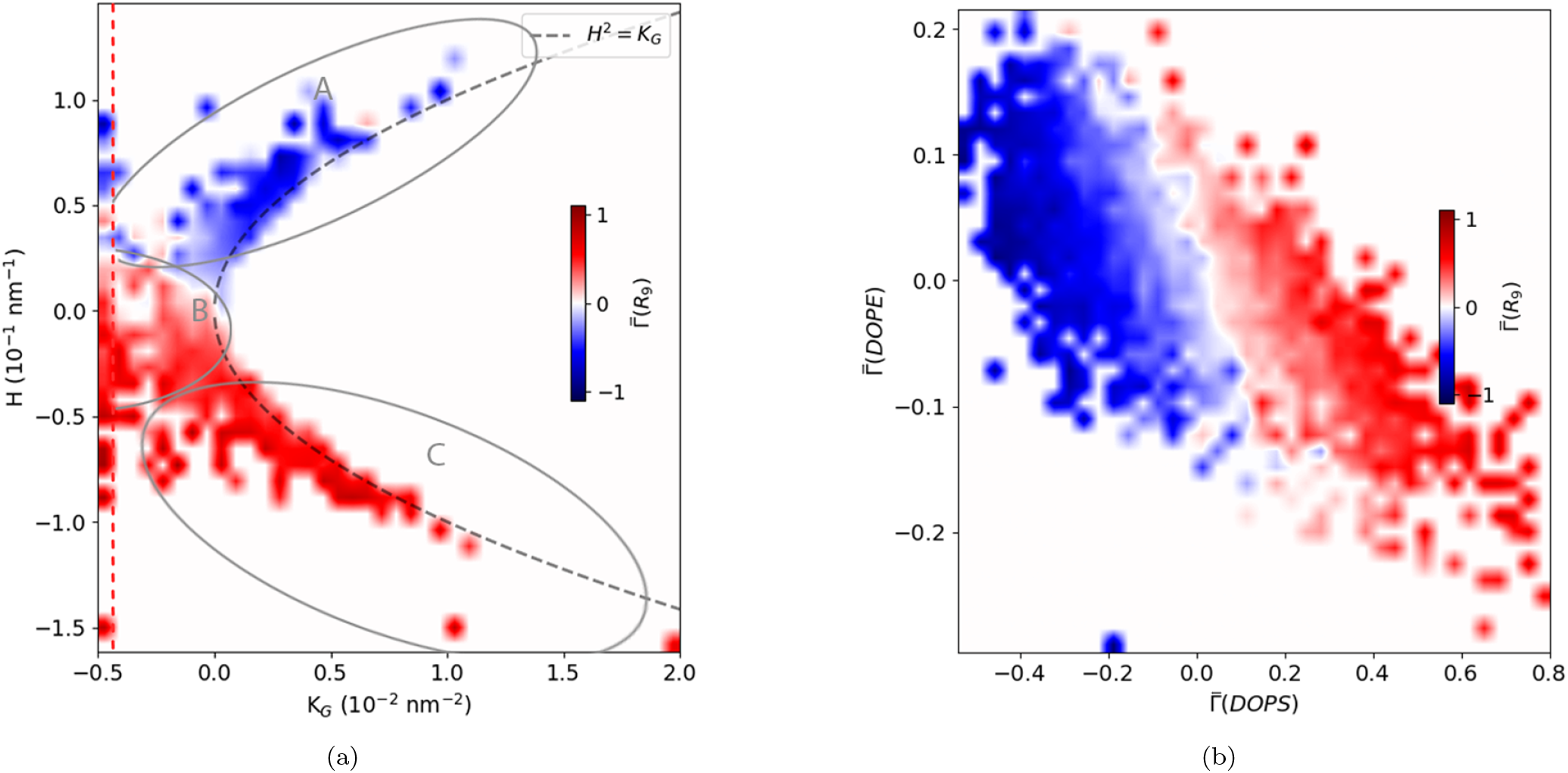
Curvature Sorting. (a) Bin-averaged heatmap of excess R_9_ coverage on stomatocyte at reduced volume (*ν*_0_) of 0.80. Regions A, B, and C correspond to the exterior, neck, and invagination, respectively. (b) Heatmap showing the excess coverage of R_9_ as a function of DOPE and DOPS composition. The dashed line delineates the excluded region by binning out all outliers to improve data clarity.

The excess coverage Γ(*R*_9_) is analyzed as a function of mean curvature (*H*) and the Gaussian curvature (*K*_*G*_) in Fig. 9a. Region A, corresponding to the vesicle exterior (*H >* 0, *K*_*G*_ *>* 0), shows a depletion of R_9_. In contrast, Region B, representing the invagination neck (*H* ∼ 0, *K*_*G*_ *<* 0), and Region C, representing the interior of the bud (*H <* 0, *K*_*G*_ *>* 0), exhibit a significant enrichment of R_9_. These observations underscore a strong dependence of R_9_ localization on membrane curvature.

This noticeable R_9_ localization at the neck and invagination regions is accompanied by significant rearrangements of DOPE and DOPS lipids, as illustrated in Fig. 9b. These lipids exhibit pronounced negative curvatures upon interaction with R_9_. Such curvature-dependent sorting illustrates the mechanistic role of lipid-peptide interactions in driving the spatial organization of R_9_ on complex membrane geometries: An obvious component of the stabilization of the inward bud is the induction of negative curvature by R_9_ on DOPS. It is natural to expect this “couple” to sort to a location where its energy is minimized. However, in our simulations, the binding energy of peptides to lipids also mediates the induced lipid sorting. R_9_ molecules have a lower mixing entropy than the lipids and cover multiple lipids by one peptide. In our model, the presence of R_9_ comes without a *per face* internal mixing entropy. Nevertheless, the expansion of R_9_ on its mesh lattice sites is associated with its own entropy, reflected by its wide distribution. This entropy emerges from generating Monte Carlo scheme, which distributes the peptides on the mesh. Yet, this “moving lipid bracket” formed by the peptide facilitates the symmetry breaking necessary for bud stabilization. This behavior, together with the curvature generation, is the mesoscopic consequence of the behavior we extracted from molecular simulations in flat geometries. Despite its enduring ability to reproduce the morphology of cell penetration, the DOPE/DOPC/DOPS model system’s effectiveness in creating these analogies remains unexplained.

## 4 Conclusion

We investigated peptide binding, lipid demixing and curvature generation of a model system for cell penetration using atomistic molecular dynamics simulations. The chemical specificity of lipid bilayer interaction with the efficient cell-penetrating peptide R_9_ over K_9_ is visible both in the binding energies and in the curvature generation. R_9_ binds more strongly and generates more *negative* curvature than K_9_ for all examined lipids. In contrast to other studies, we find this effect to be unrelated to guanidinium pairing. Instead, the deeper penetration of the guanidinium sidechains into the headgroup region of the membrane is responsible for the observed differences.

We systematically transferred these specific interactions into effective parameters of a continuum model. After validation of this model, we used it to elucidate the mesoscopic *consequences* of chemical specificity.

Inward budding is believed to be the initial step of cell penetration. ^7^ Our mesoscopic DTS simulations show how the R_9_ peptide stabilizes inward buds. These invaginations exhibit a strong tendency for R_9_ localization in regions of negative mean and negative Gaussian curvature. The formation of such structures requires sufficient available membrane, corresponding to a reduced volume of approximately 0.85, thereby requiring either an excess membrane area or the application of hyperosmotic stress to induce the transition. This poses a challenge for lipid vesicle experiments (where the surface/volume ratio would have to be modified by vesicle fusion or osmotic pressure), but not for cells with rough membrane surfaces. K_9_ was found to be unable to generate this type of structure, in agreement with its known inability to passively enter mammalian cells. Our findings indicate that strong binding and curvature generation are key to R_9_ cell penetration. The crosslinking and membrane-aggregating effect of R_9_ on membranes is the known unknown in the mechanism of CPP entry. Finally, these results do not contradict our previous investigations^3^ in any way, as multilamellarity and fusion are expected to arise in the follow-on steps.

## Author Contributions

CA designed and supervised the research. KB and CA performed MD simulations and their analysis. JK performed DTS simulations and analysis. The model was implemented by CA and JK. CA, JK and KB wrote the paper.

## Acknowledgement

CA and JK were supported by Charles University PRIMUS grant (PRIMUS/20/SCI/015). JK wishes to thank Charles University for the UNCE Math MAC Scholarship (UNCE/24/SCI/005). KB acknowledges support from Charles University, where she is enrolled as a Ph.D. student. KB acknowledges HPCg at IOCB Prague for computational resources.

## References

1. Misawa, T. Cell-Penetrating Peptides; John Wiley & Sons, Ltd, 2023; Chapter 12, pp 203–218.

2. Vazdar, M.,, Heyda, J.; Mason, P. E.; Tesei, G.; Allolio, C.; Lund, M.; Jungwirth, P. Arginine “Magic”: Guanidinium Like-Charge Ion Pairing from Aqueous Salts to Cell Penetrating Peptides. Acc. Chem. Res. 2018, 51, 1455–1464.

3. Allolio, C.; Magarkar, A.; Jurkiewicz, P.; Baxová, K.; Javanainen, M.; Mason, P. E.; Šachl, R.; Cebecauer, M.; Hof, M.; Horinek, D.; Heinz, V.; Rachel, R.; Ziegler, C. M.; Schröfel, A.; Jungwirth, P. Arginine-rich cell-penetrating peptides induce membrane multilamellarity and subsequently enter via formation of a fusion pore. Proc. Natl. Acad. Sci. USA 2018, 115, 11923–11928.

4. Mishra, A.; Lai, G. H.; Schmidt, N. W.; Sun, V. Z.; Rodriguez, A. R.; Tong, R.; Tang, L.; Cheng, J.; Deming, T. J.; Kamei, D. T.; Wong, G. C. L. Translocation of HIV TAT peptide and analogues induced by multiplexed membrane and cytoskeletal interactions. Proc. Natl. Acad. Sci. USA 2011, 108, 16883–16888.

5. Schmidt, N.; Mishra, A.; Lai, G. H.; Wong, G. C. Arginine-rich cell-penetrating peptides. FEBS Lett. 2010, 584, 1806– 1813.

6. Hirose, H.; Takeuchi, T.; Osakada, H.; Pujals, S.; Katayama, S.; Nakase, I.; Kobayashi, S.; Haraguchi, T.; Futaki, S. Transient Focal Membrane Deformation Induced by Arginine-rich Peptides Leads to Their Direct Penetration into Cells. Mol. Therapy 2012, 20, 984–993.

7. Sahni, A.; Ritchey, J. L.; Qian, Z.; Pei, D. Cell-Penetrating Peptides Translocate across the Plasma Membrane by Inducing Vesicle Budding and Collapse. J. Am. Chem. Soc. 2024, 146, 25371–25382.

8. Fuchs, S. M.; Raines, R. T. Pathway for Polyarginine Entry into Mammalian Cells. Biochemistry 2004, 43, 2438–2444.

9. Säalik, P.; Niinep, A.; Pae, J.; Hansen, M.; Lubenets, D.; Ülo Langel Pooga, M. Penetration without cells: Membrane translocation of cell-penetrating peptides in the model giant plasma membrane vesicles. J. Control. Release 2011, 153, 117–125.

10. Allolio, C.; Baxova, K.; Vazdar, M.; Jungwirth, P. Guanidinium Pairing Facilitates Membrane Translocation. J. Phys. Chem. B 2016, 120, 143–153.

11. Robison, A. D.; Sun, S.; Poyton, M. F.; Johnson, G. A.; Pellois, J.-P.; Jungwirth, P.; Vazdar, M.; Cremer, P. S. Polyarginine Interacts More Strongly and Cooperatively than Polylysine with Phospholipid Bilayers. J. Phys. Chem. B 2016, 120, 9287–9296.

12. Vazdar, M.; Wernersson, E.; Khabiri, M.; Cwiklik, L.; Jurkiewicz, P.; Hof, M.; Mann, E.; Kolusheva, S.; Jelinek, R.; Jungwirth, P. Aggregation of Oligoarginines at Phospholipid Membranes: Molecular Dynamics Simulations, Time-Dependent Fluorescence Shift, and Biomimetic Colorimetric Assays. J. Phys. Chem. B 2013, 117, 11530–11540.

13. Nguyen, M. T. H.; Biriukov, D.; Tempra, C.; Baxova, K.; Martinez-Seara, H.; Evci, H.; Singh, V.; Šachl, R.; Hof, M.; Jungwirth, P.; Javanainen, M.; Vazdar, M. Ionic Strength and Solution Composition Dictate the Adsorption of Cell-Penetrating Peptides onto Phosphatidylcholine Membranes. Langmuir 2022, 38, 11284–11295.

14. Lamazière, A.; Burlina, F.; Wolf, C.; Chassaing, G.; Trugnan, G.; Ayala-Sanmartin, J. Non-Metabolic Membrane Tubulation and Permeability Induced by Bioactive Peptides. PLOS ONE 2007, 2, 1–11.

15. Sahni, A.; Qian, Z.; Pei, D. Cell-Penetrating Peptides Escape the Endosome by Inducing Vesicle Budding and Collapse. ACS Chem. Biol. 2020, 15, 2485–2492.

16. Dougherty, P. G.; Sahni, A.; Pei, D. Understanding Cell Penetration of Cyclic Peptides. Chem. Rev. 2019, 119, 10241–10287.

17. Ramakrishnan, N.; Bradley, R. P.; Tourdot, R. W.; Radhakrishnan, R. Biophysics of membrane curvature remodeling at molecular and mesoscopic lengthscales. J. Phys.: Condens. Matter 2018, 30, 273001.

18. Leibler, S. Curvature instability in membranes. J. Phys. (France) 1986, 47, 507– 516.

19. Denisov, G.; Wanaski, S.; Luan, P.; Glaser, M.; McLaughlin, S. Binding of basic peptides to membranes produces lateral domains enriched in the acidic lipids phosphatidylserine and phosphatidylinositol 4,5-bisphosphate: An electrostatic model and experimental results. Biophys. J. 1998, 74, 731 – 744.

20. Pezeshkian, W.; Gao, H.; Arumugam, S.; Becken, U.; Bassereau, P.; Florent, J.-C.; Ipsen, J. H.; Johannes, L.; Shillcock, J. C. Mechanism of Shiga Toxin Clustering on Membranes. ACS Nano 2017, 11, 314–324.

21. Ruhoff, V. T.; Bendix, P. M.; Pezeshkian, W. Close, but not too close: a mesoscopic description of (a)symmetry and membrane shaping mechanisms. Emerg. Top. Life Sci. 2023, 7, 81–93.

22. Campelo, F.; McMahon, H. T.; Kozlov, M. M. The Hydrophobic Insertion Mechanism of Membrane Curvature Generation by Proteins. Biophys. J. 2008, 95, 2325 – 2339.

23. Noguchi, H.; Fournier, J.-B. Membrane structure formation induced by two types of banana-shaped proteins. Soft Matter 2017, 13, 4099–4111.

24. Zimmerberg, J.; Kozlov, M. M. How proteins produce cellular membrane curvature. Nat. Rev. Mol. Cell Biol. 2006, 7, 9–19.

25. Konar, S.; Arif, H.; Allolio, C. Mitochondrial membrane model: Lipids, elastic properties, and the changing curvature of cardiolipin. Biophys. J. 2023, 122, 4274– 4287.

26. Schachter, I.; Allolio, C.; Khelashvili, G.; Harries, D. Confinement in Nanodiscs Anisotropically Modifies Lipid Bilayer Elastic Properties. J. Phys. Chem. B 2020, 124, 7166–7175.

27. Allolio, C.; Harries, D. Calcium Ions Promote Membrane Fusion by Forming Negative-Curvature Inducing Clusters on Specific Anionic Lipids. ACS Nano 2021, 15, 12880–12887.

28. Starke, L. J.; Allolio, C.; Hub, J. S. How pore formation in complex biological membranes is governed by lipid composition, mechanics, and lateral sorting. PNAS Nexus 2025, pgaf033.

29. Pelech, P.; Navarro, P. P.; Vettiger, A.; Chao, L. H.; Allolio, C. Stress-mediated growth determines E. coli division site morphogenesis. bioRxiv 2024, 2024.09.11.612282.

30. Sun, D.; Forsman, J.; Woodward, C. E. Atomistic Molecular Simulations Suggest a Kinetic Model for Membrane Translocation by Arginine-Rich Peptides. J. Phys. Chem. B 2015, 119, 14413–14420.

31. Allolio, C.; Fábián, B.; Dostalík, M. OrganL: Dynamic triangulation of biomembranes using curved elements. Biophys. J. 2024, 123, 1553–1562.

32. Lee, J.; Patel, D. S.; Ståhle, J.; Park, S.-J.; Kern, N. R.; Kim, S.; Lee, J.; Cheng, X.; Valvano, M. A.; Holst, O.; Knirel, Y. A.; Qi, Y.; Jo, S.; Klauda, J. B.; Widmalm, G.; Im, W. CHARMM-GUI Membrane Builder for Complex Biological Membrane Simulations with Glycolipids and Lipoglycans. J. Chem. Theory Comput. 2019, 15, 775–786.

33. Nencini, R.; Tempra, C.; Biriukov, D.; Polák, J.; Ondo, D.; Heyda, J.; Ollila, S.; Javanainen, M.; Martinez-Seara, H. Prosecco: polarization reintroduced by optimal scaling of electronic continuum correction origin in MD simulations. Biophys. J. 2022, 121, 157a.

34. Abraham, M. J.; Murtola, T.; Schulz, R.; Páll, S.; Smith, J. C.; Hess, B.; Lindahl, E. GROMACS: High Performance Molecular Simulations through Multi-Level Parallelism from Laptops to Supercomputers. SoftwareX 2015, 1–2, 19 – 25.

35. Nosé, S. A Molecular Dynamics Method for Simulations in the Canonical Ensemble. Mol. Phys. 1984, 5, 255–268.

36. Hoover, W. G. Canonical Dynamics: Equilibrium Phase-Space Distributions. Phys. Rev. A 1985, 31, 1695–1697.

37. Parrinello, M.; Rahman, A. Crystal Structure and Pair Potentials: A Molecular-Dynamics Study. Phys. Rev. Lett. 1980, 45, 1196.

38. Essman, U.; Perela, L.; Berkowitz, M. L.; Darden, T.; Lee, H.; Pedersen, L. G. A Smooth Particle Mesh Ewald Method. J. Chem. Phys. 1995, 103, 8577–8593.

39. Hess, B.; Bekker, H.; Berendsen, H. J. C.; Fraaije, J. G. E. M. LINCS: A Linear Constraint Solver for Molecular Simulations. J. Comp. Chem. 1997, 18, 1463–1472.

40. Miyamoto, S.; Kollman, P. A. Settle: An Analytical Version of the SHAKE and RATTLE Algorithm for Rigid Water Models. J. Comp. Chem. 1992, 13, 952–962.

41. Sega, M.; Fábian, B.; Jedlovszky, P. Pressure Profile Calculation with Mesh Ewald Methods. J. Chem. Theory Comput. 2016, 12, 4509–4515.

42. Allolio, C.; Haluts, A.; Harries, D. A Local Instantaneous Surface Method for Extracting Membrane Elastic Moduli from Simulation: Comparison with other Strategies. Chem. Phys. 2018, 514, 31 – 43.

43. Rawicz, W.; Olbrich, K.; McIntosh, T.; Needham, D.; Evans, E. Effect of Chain Length and Unsaturation on Elasticity of Lipid Bilayers. Biophys. J. 2000, 79, 328– 339.

44. Binder, H.; Gawrisch, K. Effect of Unsaturated Lipid Chains on Dimensions, Molecular Order and Hydration of Membranes. J. Phys. Chem. B 2001, 105, 12378–12390.

45. Arun, K. S.; Huang, T. S.; Blostein, S. D. Least-Squares Fitting of Two 3-D Point Sets. IEEE Trans. Pattern Anal. Mach. Intell 1987, PAMI-9, 698–700.

46. Biriukov, D.; Osifova, Z.; Nguyen, M.; Mason, P.; Dračínský, M.; Jungwirth, P.; Heyda, J.; Morandi, M.; Vazdar, M. The Origins of Arginine “Magic”: Guanidinium Like-Charge Ion Pairing and Oligoarginine Aggregation in Water by NMR, Cryoelectron Microscopy, and Molecular Dynamics Simulations. bioRxiv 2024, 2024.08.04.606526.

47. Khelashvili, G.; Weinstein, H.; Harries, D. Protein Diffusion on Charged Membranes: A Dynamic Mean-Field Model Describes Time Evolution and Lipid Reorganization. Biophys. J. 2008, 94, 2580–2597.

48. Khelashvili, G.; Harries, D.; Weinstein, H. Modeling Membrane Deformations and Lipid Demixing upon Protein-Membrane Interaction: The {BAR} Dimer Adsorption. Biophys. J. 2009, 97, 1626 – 1635.

49. Szleifer, I.; Kramer, D.; Ben-Shaul, A.; Gelbart, W. M.; Safran, S. A. Molecular theory of curvature elasticity in surfactant films. J. Chem. Phys. 1990, 92, 6800–6817.

50. Hamm, M.; Kozlov, M. Elastic Energy of Tilt and Bending of Fluid Membranes. Eur. Phys. J. E 2000, 3, 323–335.

51. Helfrich, W. Elastic Properties of Lipid Bilayers: Theory and Possible Experiments. Z. Naturforsch. C 1973, 28, 693–793.

52. Canham, P. The minimum energy of bending as a possible explanation of the biconcave shape of the human red blood cell. J. Theor. Biol. 1970, 26, 61–81.

53. Evans, E. Bending resistance and chemically induced moments in membrane bilayers. Biophys. J. 1974, 14, 923–931.

54. Linke, G. T.; Lipowsky, R.; Gruhn, T. Osmotically induced passage of vesicles through narrow pores. Europhys. Lett. 2006, 74, 916.

55. Khelashvili, G.; Kollmitzer, B.; Heftberger, P.; Pabst, G.; Harries, D. Calculating the Bending Modulus for Multicomponent Lipid Membranes in Different Thermodynamic Phases. J. Chem. Theory Comput. 2013, 9, 3866–3871.

56. Seifert, U.; Berndl, K.; Lipowsky, R. Shape transformations of vesicles: Phase diagram for spontaneous-curvature and bilayer-coupling models. Phys. Rev. A 1991, 44, 1182–1202.

57. Hossein, A.; Deserno, M. Spontaneous Curvature, Differential Stress, and Bending Modulus of Asymmetric Lipid Membranes. Biophys. J. 2020, 118, 624–642.

58. Hu, M.; De Jong, D.; Marrink, S.; Deserno, M. Gaussian curvature elasticity determined from global shape transformations and local stress distributions: A comparative study using the MARTINI model. Faraday Discuss. 2012, 161, 365–382.

59. Bian, X.; Litvinov, S.; Koumoutsakos, P. Bending models of lipid bilayer membranes: Spontaneous curvature and area-difference elasticity. Comput. Methods Appl. Mech. Eng. 2020, 359, 112758.

60. Derganc, J. Curvature-driven lateral segregation of membrane constituents in Golgi cisternae. Phys. Biol. 2007, 4, 317.

61. Callan-Jones, A.; Sorre, B.; Bassereau, P. Curvature-driven lipid sorting in biomembranes. Cold Spring Harb. Perspect. Biol. 2011, 3, a004648–a004648.

