## Supporting Information for "From Molecular Insight to Mesoscale Membrane Remodeling: Curvature Generation by Arginine-Rich Cell-Penetrating Peptides"

### Supplementary to Multiscale Modelling and Chemical Specificity of Lipid Membranes; Case of Arginine Magic

April 14, 2025

#### 1 Parameter Extraction from MD

##### 1.1 Stress Tensor

The local stress tensor was calculated using the rerun functionality of the gmx mdrun module of our implementation[1] of a Goetz-Lipowsky-Decomposition[2] into the code by Sega *et al.*[3] The modified version of GROMACS is available at `https://github.com/allolio`. The same cutoffs were applied as described above. Our approach is similar to the method in Ref. [4]. Specifically, we set the normal pressure $p_{\mathrm{N}}$ to impose zero surface tension:

$$\sigma = \int_{-l}^{l} \pi(z)\,\mathrm{d}z = \int_{-l}^{l} \Big[ -\frac{1}{2}\{p_{xx}(z) + p_{yy}(z)\} + p_{\mathrm{N}} \Big]\,\mathrm{d}z = 0. \tag{1}$$

Here, $\sigma$ is obtained from the diagonal components of the stress tensor [5], and the integral is carried out over the entire simulation box $[-l, l]$ along the membrane normal $z$, where $z = 0$ corresponds to the bilayer center. Since this approach requires $p_{\mathrm{N}}$ being close to 1 bar, we validated that the deviations from this target were $\leq 3.2$ bar for all simulations.

##### 1.2 Extraction of Model Parameters

The bending modulus $\kappa$ and the tilt modulus $\kappa_\theta$ were calculated using the Re-SIS method.[6] We used lipid director definitions and an implementation of the LIPIDATOR-TOOLKIT available at `https://github.com/allolio/lipidator-toolkit`. This allowed the use of the well-known[7, 8] relation of the first bending moment to the product of spontaneous curvature, $J_{\mathrm{s}}$ and $\kappa$:

$$\kappa J_{\mathrm{s}} = \int_{0}^{l} \pi(z) z \mathrm{d}z. \tag{2}$$

Here, integration is carried out along the membrane normal $\hat{\mathbf{e}}_z$ and only over one monolayer, and, therefore, monolayer elastic properties are used. The reported values are averages over both monolayers. The bilayer values can be obtained by summing up the monolayer values, resulting in a bilayer $J_{\mathrm{s}}^{\mathrm{b}} = 0$ for symmetric membranes.

##### 1.2.1 Excess Lipid Number

The number of lipids bound to peptide was extracted from the scaled RDF data from MD using the 2D Kirkwood Buff Integral:

$$\Gamma = \sigma_\beta \int 2\pi r(g_{\alpha\beta}(r) - 1)\mathrm{d}r, \tag{3}$$

where $\sigma_\beta$ is the surface density of $\beta$-particles. The RDF values used are the averages of "Charmm" and "Prosecco".

#### 1.3 Binding Energies

The binding energies were computed via Umbrella sampling at 300 K, the setup used 128 lipids (either DOPE, DOPC, or DOPS), 14 K $H_2O$ and 150 mM ions, in addition to neutralizing counterions. This corresponds to 34 $K^+$ and 43 $Cl^-$ for neutral membranes and 153 $K^+$, 34 $Cl^-$ for DOPS. Each umbrella window was run for 250 ns after pre-equilibration. The pull coordinate was the center of mass distance along the membrane normal, between lipids and the peptides. The force constant was typically 1000 kJ/mol/nm$^2$, and 21 windows were used for each simulation. The binding energies were computed using a charge scaling derivative of CHARMM36m/CHARMM36 force field - ProsECCo75[9] with the TIP3P water model.

### 2 The Mesoscopic Model

The simulations use the OrganL package.[10] We want to describe how $R_9$ (and $K_9$) binds to bilayers composed of mixed lipids and influences the membrane intrinsic curvature $J_S$ as well as bending rigidity $\kappa$, as defined in the Canham-Helfrich[11, 12] free energy

$$F_{\mathrm{HF}} = \int_{\mathrm{Surf}} \mathrm{d}A \left\{ \frac{\kappa}{2}(2H - J_S)^2 \right\}. \tag{4}$$

For this purpose, we construct a model describing the curvature influence for arbitrary mixtures (as we assume that lipid demixing occurs). In other words, $\kappa$ and $J_S$ can vary locally. We also want to simulate the vesicle geometries under realistic conditions of osmotic pressure and surface tension. The protein binding model closely resembles that of a **regular solution**, i.e. we will assume the ideal mixing of lipids (a standard assumption) together with a fixed, incremental binding energy of the peptide up to the maximum concentration. To support the latter assumption, we note that according to Cremer et al.[13], the binding of $R_9$ in particular is noncooperative, i.e., modeled well by a Langmuir isotherm, which implies noninteracting binding sites.

#### 2.1 Effect of Lipids

##### 2.1.1 Monolayer

We calculate the bending modulus as a harmonic average of the single lipid moduli $\kappa_i$:

$$\frac{1}{\kappa} = \sum_i \phi_i \frac{1}{\kappa_i}. \tag{5}$$

Here, $\phi_i$ refers to the volume fraction, which is calculated as

$$\phi_i = \frac{m_i A_i}{\sum_j m_j A_j}, \tag{6}$$

with $A_i$ denoting the area per lipid of the lipid species $i$. We assume that the area per lipid does not change while mixing. The area per lipid $A_i$ is a reference area contributing to $A_0$ so that mixing/bending and stretching are not coupled. By $m_i$, we refer to the number of lipids of species $i$ present in a triangle. Of course, a completely different behaviour is possible. An example occurs during a phase transition to the gel or the $L_O$ phase. As long as the behavior is well-defined, it is easy to implement the requisite models in our code.

The spontaneous curvature is computed as

$$J_s = \sum_i \phi_i c^0_{s,i}, \tag{7}$$

where $c^0_i$ is spontaneous curvature of the pure $i$.[14, 15]We have recently tested the validity of these equations against atomistic molecular dynamics results.[4] Furthermore, the mixing entropy of the lipid composition with a total occupancy of lipids $M$ on a side/leaflet of a triangle can be estimated as ideal in the surface fractions[14]:

$$F_{\text{mix}} = kTM \sum_i \phi_i \ln \phi_i. \tag{8}$$

##### 2.1.2 Bilayer

For the averaging of the two sides, we use the indices $u, d$ for the upper and lower monolayers, respectively, and $b$ for the bilayer:

$$\begin{aligned} \kappa_b &= \kappa_u + \kappa_b; \\ F_{\text{mix,b}} &= F_{\text{mix,u}} + F_{\text{mix,d}} \end{aligned} \tag{9}$$

The spontaneous curvature is computed as:

$$J_{\text{s,b}} = \frac{1}{2}\big(J_{s,u} - J_{s,d}\big). \tag{10}$$

The sign change is due to the change in a normal orientation and curvature direction when flipping between two faces. The resulting equations are equivalent to averaging over two monolayers, up to a constant offset, under the condition that $\kappa_u \approx \kappa_d$ and the thickness $d \to 0$. The error bars for $\kappa$ and $\kappa^b_\theta$ were obtained by Gaussian fitting to the tilt and the divergence of the tilt distribution, respectively. Ten separate distributions were computed by sampling independent parts of the simulation trajectory; the standard deviations from values thus obtained were used to calculate the standard error over the mean of the entire trajectory. In the case of $J_s$, the monolayers were averaged, and three windows were used to compute the standard error for $J_S\kappa$. We computed Gaussian error propagation to obtain the error bars.

#### 2.2 Effect of Peptides

The peptides are assumed to occupy an area $A_p$, *on* a lipid monolayer. Then, the coverage of the peptide is computed by

$$\phi_p = n_p \frac{A_p}{A_l}, \tag{11}$$

where $n_p$ is the number of peptides. $\phi_p$ is controlled not to exceed 1. $A_l$ is the total area of the lipid patch, i.e.

$$A_l = \sum m_i A_i, \tag{12}$$

which is calculated on a leaflet. For each pure lipid $i$, there is a binding free energy $\mu_i$ with the peptide. Accordingly, the total binding energy of the peptide is estimated as

$$F_b = n_p \sum_i \phi_i \mu_i. \tag{13}$$

The influence of the peptide on the lipid membrane properties is further evaluated by analyzing the lipid surface fractions within the membrane patch. Each pure lipid species interacting with the peptide contributes its respective bending rigidity, $\kappa_i$, which differs from the value observed in the absence of the peptide. The net value is computed as:

$$\frac{1}{\kappa_p} = \sum_i \phi_i \frac{1}{\kappa_i} \tag{14}$$

and finally weighted as

$$\frac{1}{\kappa} = \phi_p \frac{1}{\kappa_p} + (1 - \phi_p) \frac{1}{\kappa_n}, \tag{15}$$

where $\kappa_n$ is the bending rigidity computed without peptide influence as in the previous section.

Similarly,

$$J_{s,p} = \sum_i \phi_i c0_i \tag{16}$$

and

$$J_s = \phi_p J_{\mathrm{s,p}} + (1 - \phi_p) J_{\mathrm{s,n}}. \tag{17}$$

Note that the peptide does not contribute to the mixing entropy, as it is considered large with respect to the mixture. The number of peptides is always an integer, but the number of lipids can be rational number on each face.

#### 2.3 Gaussian Curvature

In the absence of a good estimator for the Gaussian curvature, the bilayer Gaussian curvature is computed as[1, 16]

$$\bar{\kappa}_b = -0.8\kappa - 4 J_s \kappa d, \tag{18}$$

which is a result of an expansion to linear order in the membrane thickness from bilayer midplane $d$, taken to be 1.2 nm. The prefactor of 0.8 was taken by comparing monolayer Gaussian curvature moduli measured from the literature[1].

#### 2.4 Total Energy

The total energy of the system is then given by

$$F = \int_{\text{Surf.}} \mathrm{d}A \left\{ \frac{\kappa_b}{2}(2\tilde{H} - J_{S,b})^2 + \bar{\kappa}_b K_G \right\} + E_{\text{pV}} + E_{\text{stretch}} + \sum_{\text{face}} \left[ F_b + F_{\text{mix,b}} \right] \tag{19}$$

In this equation, the parameters $\kappa, \bar{\kappa}$ and $J_S$ are constant on each face as far as integration is concerned. Integration is performed over the faces. $F_b$ and $F_{\text{mix,b}}$ are computed per face and summed up. $E_{\text{pV}}$ and $E_{\text{stretch}}$ are computed using global values for $(A, A_0)$ and $(V, V_0)$, which are the (current, initial) values, respectively.

##### 2.4.1 Area and Volume Control

The film area is controlled via the area compressibility $K_A$, and the following stretching energy is added to the total system:

$$E_{\text{stretch}} = \frac{K_A}{2} \frac{(A - A_0)^2}{A_0} \tag{20}$$

We use the isothermal ideal gas equation of state to relate osmotic pressure to the vesicle volume

$$\Delta p = \Delta c R T. \tag{21}$$

Assuming that the vesicle is in equilibrium with the external pressure, the isothermal volume work, relative to the equilibrium volume $V_0$, which defines the reference state where $E_{\text{pV}} = 0$, can be expressed as

$$E_{\text{pV}} = c_0 k_B T \big[ V - V_0 - V_0 \log(|V/V_0|) \big] \tag{22}$$

where $c_0$ is concentration, $V_0$ is the initial volume, $k_B$ is the Boltzmann constant, and $T$ is the absolute temperature.

#### 2.5 Validation Details

The optimization procedure consists of two sequential sub-problems:

1. **Free Energy Minimization:** The free energy functional, defined in Eq. (5), is minimized subject to the constraints specified in Eqs. (6–8). This constrained optimization is carried out using the `trust-constr` algorithm (default `SLSQP` doesn't converge for $K_9$) from the `SciPy` optimization library, which supports the inclusion of `LinearConstraint` objects.

2. **Fitting to Simulation Data:** The above minimization is wrapped using the $\mathcal{L}^2$ norm of the discrepancy between the predicted excess quantities and those obtained from molecular dynamics (MD) simulations (as described in the main text) and is minimized. This step is implemented using the `Powell` method available in `SciPy`.

Table 1: Summary of model parameters for lipid bilayers with/without peptides[1].

| no peptide | $\kappa$ [$kT$] | $c^0$ [$nm^{-1}$] | $A_0$ [nm] | $E_{bind}$ [kJ/mol] |
|---|---|---|---|---|
| DOPE | 15.83 | -0.24 | 0.625 | $\times$ |
| DOPS | 14.27 | -0.09 | 0.643 | $\times$ |
| DOPC | 11.56 | 0.0 | 0.679 | $\times$ |

(a) No peptide.

| $R_9$ | $\kappa$ [$kT$] | $c^0$ [$nm^{-1}$] | $A_0$ [nm] | $E_{bind}$ [kJ/mol] |
|---|---|---|---|---|
| DOPE | 16.25 | $-0.30$ | 0.625 | $-23.2 \pm 2.6$ |
| DOPS | 13.60 | $-0.25$ | 0.643 | $-86.0 \pm 4.3$ |
| DOPC | 10.88 | $-0.04$ | 0.679 | $-19.6 \pm 3.0$ |

(b) With $R_9$.

| $K_9$ | $\kappa$ [$kT$] | $c^0$ [$nm^{-1}$] | $A_0$ [nm] | $E_{bind}$ [kJ/mol] |
|---|---|---|---|---|
| DOPE | 15.83 | $-0.24$ | 0.625 | $-0.7 \pm 1.3$ |
| DOPS | 16.26 | $-0.17$ | 0.643 | $-51.4 \pm 2.3$ |
| DOPC | 11.56 | 0.0 | 0.679 | $\times$ |

(c) With $K_9$.

#### 2.6 Mesoscopic Monte Carlo Simulations

The model parameters used for the simulations are summarized in Table 1.

The starting point for the simulation was an equilateral spherical mesh of radius 50 nm with 2552 faces (so that the area per face matches the (R9) area).

A stable stomatocyte geometry was generated from the spherical mesh in $\approx 3 \times 10^5$ MC steps (with step size of 1.0 for both `VertexMove` and `NormalMove`; and `autoTune` of 10 steps) [10] by setting the reduced volume

$$\nu_0 := \frac{6V}{(A)^{3/2}}\sqrt{\pi} = \sqrt{2}^{-1}. \tag{23}$$

This corresponds to the situation of two vesicles of equal size having fused, i.e. doubling $A$ and $V$ from an initial spherical geometry ($\nu_0 = 1$). This was achieved by reducing $V$ while keeping $A$ constant. To get a stomatocyte structure with a good constriction at the neck, it is necessary to simulate back and forth by changing the values of $V_0$ at $\sim 0.85$, with high `Remesh` values [10](controls how often the bond flipping move is triggered in OrganL). This is essential to get a minimal number of faces at the neck region (as necessary for constriction). This stable geometry is used as a starting point for all the other stomatocyte simulations (control, $R_9$ and $K_9$).

To scan the energy profile as a function of $\nu_0$, we gradually increase $V$ back to the initial value (corresponding to the sphere). Subsequently, each $V$ value was sampled until the standard deviation in energy was minimized, ensuring convergence (at least $2 \times 10^6$ MC steps).

#### 2.7 Analysis

- Energies are computed by averaging the final 500 outputs, corresponding to the last $500,000$ Monte Carlo steps, following an initial equilibration phase of at least $2 \times 10^6$ MC steps from the starting geometry.

- A *mesh averaging* procedure is applied purely for visualization purposes using the Iterative Closest Point (ICP) algorithm [17] over the last 200 output files. The face properties are averaged and mapped onto this ICP-generated mesh to produce the representative images included in the manuscript.

- To preserve spatial correlations, a *bin-averaging* approach is employed on the last 200 outputs (spanning 200 k MC steps). In this method, Property I from each snapshot is binned according to the values of Property II, and the binned values are subsequently averaged across all snapshots. This procedure ensures the retention of correlations between Property I and Property II, facilitating the analysis of effects such as mean and Gaussian curvature variations or lipid demixing in the presence of peptides, as discussed in the manuscript.

#### 2.8 Results

Fig. 1 presents the full energy decomposition, provided as supplementary information to complement the main plot in the primary manuscript (Figure 8).

Further, for the sake of completeness, bin-averaged heating of the sorting mechanism on unfrozen meshes is plotted in Fig. 2. Comparing with the frozen case in the manuscript, it may be observed that this is a very robust but not as pronounced, owing to the inclusion of mesh fluctuation.

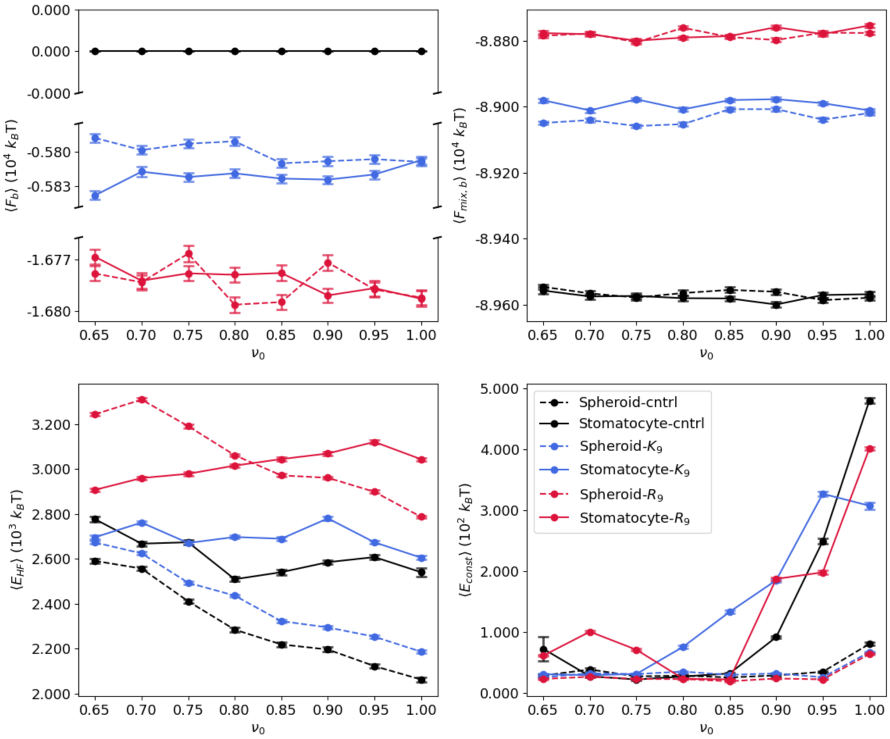


Figure 1: **Energy Decomposition Plot** Energy profiles of spheroid and stomatocyte morphologies in DOPE/DOPC/DOPS vesicles are presented under three conditions: (a) without peptides, (b) with $K_9$, and (c) in the presence of $R_9$. Error bars represent the SEM with 95% CI.

### References


[1] Christoph Allolio and Daniel Harries, *ACS Nano*, **2021**, *15*(8), 12880–12887.

[2] Rüdiger Goetz and Reinhard Lipowsky, *J. Chem. Phys.*, **1998**, *108*(17), 7397–7409.

[3] Marcello Sega, Balázs Fábián, and Pál Jedlovszky, *J. Chem. Theory Comput.*, **2016**, *12*(9), 4509–4515.

[4] Sukanya Konar, Hina Arif, and Christoph Allolio, *Biophys. J.*, **2023**, *122*(21), 4274–4287.

[5] P. Schofield, James R. Henderson, and John Shipley Rowlinson, *Proc. R. Soc. A*, **1982**, *379*(1776), 231–246.

[6] Christoph Allolio, Amir Haluts, and Daniel Harries, *Chem. Phys.*, **2018**, *514*, 31 – 43.

[7] Igal Szleifer, Diego Kramer, Avinoam Ben-Shaul, William M. Gelbart, and S. A. Safran, *J. Chem. Phys.*, **1990**, *92*(11), 6800–6817.

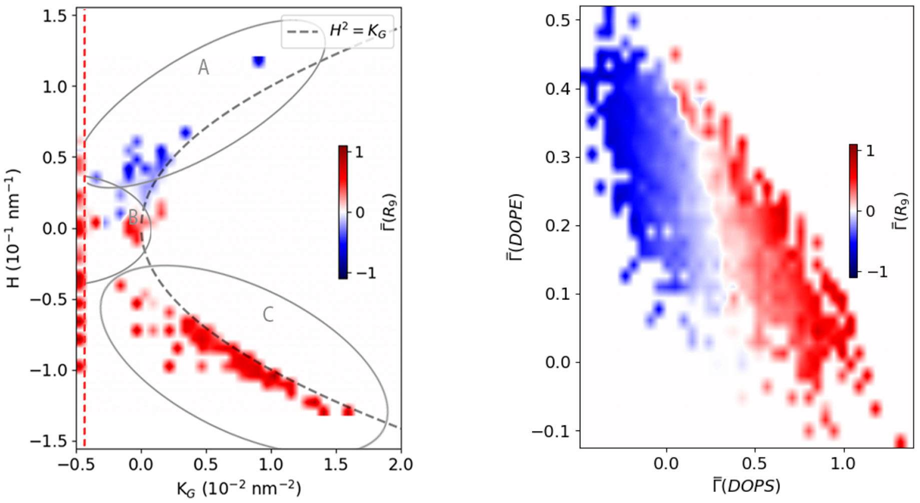


(a) (b)

Figure 2: **Curvature Sorting on unfrozen mesh.** (a) Bin-averaged heatmap of excess $R_9$ coverage on stomatocyte at reduced volume ($\nu_0$) of 0.80. Regions A, B, and C correspond to the exterior, neck, and invagination, respectively. (b) Heatmap showing the excess coverage of $R_9$ as a function of DOPE and DOPS composition. The dashed line delineates the excluded region by binning out all outliers to improve data clarity.


[8] M. Hamm and M.M. Kozlov, *Eur. Phys. J. E*, **2000**, *3*(4), 323–335.

[9] Ricky Nencini, Carmelo Tempra, Denys Biriukov, Jakub Polák, Daniel Ondo, Jan Heyda, Samuli Ollila, Matti Javanainen, and Hector Martinez-Seara, *Biophys. J.*, **2022**, *121*, 157a.

[10] Christoph Allolio, Balázs Fábián, and Mark Dostalík, *Biophys. J.*, **2024**, *123*(12), 1553–1562.

[11] W. Helfrich, *Z. Naturforsch. C*, **1973**, *28*, 693–793.

[12] P.B. Canham, *J. Theor. Biol.*, **1970**, *26*(1), 61–81.

[13] Aaron D. Robison, Simou Sun, Matthew F. Poyton, Gregory A. Johnson, Jean-Philippe Pellois, Pavel Jungwirth, Mario Vazdar, and Paul S. Cremer, *J. Phys. Chem. B*, **2016**, *120*(35), 9287–9296.

[14] D. Andelman, M. M. Kozlov, and W. Helfrich, *Europhys. Lett.*, **1994**, *25*(3), 231.

[15] George Khelashvili, Daniel Harries, and Harel Weinstein, *Biophys. J.*, **2009**, *97*(6), 1626 – 1635.

[16] Mingyang Hu, John J Briguglio, and Markus Deserno, *Biophys. J.*, **2012**, *102*(6).

[17] K. S. Arun, T. S. Huang, and S. D. Blostein, *IEEE Trans. Pattern Anal. Mach. Intell*, **1987**, *PAMI-9*(5), 698–700.
